# Short-term assessment of subfoveal injection of AAV2-*hCHM* gene augmentation in choroideremia using adaptive optics ophthalmoscopy

**DOI:** 10.1101/2021.09.17.459817

**Authors:** Jessica I. W. Morgan, Yu You Jiang, Grace K. Vergilio, Leona W. Serrano, Denise J. Pearson, Jean Bennett, Albert M. Maguire, Tomas S. Aleman

## Abstract

Subretinal injection for gene augmentation in retinal degenerations forcefully detaches the neural retina from the retinal pigment epithelium (RPE), potentially damaging photoreceptors and/or RPE cells. Here, we use adaptive optics scanning light ophthalmoscopy (AOSLO) to assess the short-term integrity of the cone mosaic following subretinal injections of AAV2-*hCHM* gene augmentation in subjects with choroideremia (CHM). Nine adult CHM patients received uniocular subfoveal injections of low dose (5x10^10^ vector genome (vg) per eye, n=5) or high dose (1x10^11^ vg per eye, n=4) AAV2-*hCHM*. The macular regions of both eyes were imaged pre- and one-month post-injection using a custom-built, multimodal AOSLO. Post-injection cone inner segment mosaics were compared to pre-injection mosaics at multiple regions of interest (ROIs). Post-injection AOSLO images showed preservation of the cone mosaic in all 9 AAV2-*hCHM* injected eyes. Mosaics appeared intact and contiguous one-month post-injection, with the exception of foveal disruption in one patient. Co-localized optical coherence tomography showed foveal cone outer segment (COS) shortening post-injection (significant, n=4; non-significant, n=4; unchanged, n=1). Integrity of the cone mosaic is maintained following subretinal delivery of AAV2-*hCHM*, providing strong evidence in support of the safety of the injections. Minor foveal thinning observed following surgery corresponds with short-term COS shortening rather than cone cell loss.

## Introduction

The advent of gene therapy for inherited retinal degenerations has revolutionized the field of ophthalmic care.^1^ The FDA’s recent approval of LUXTURNA® has provided the first clinical available treatment option for RPE65 associated inherited retinal degeneration^2^ and given hope to numerous other patients who suffer from genetic blinding disease without available treatments.^3^ Indeed, numerous clinical trials testing gene augmentation to treat other inherited retinal degenerations (IRDs) are being conceived or are in progress.^4–6^

Choroideremia (CHM) is one such IRD where gene augmentation is being tested in multi-institutional gene therapy clinical trials.^7–18^ CHM is an X-linked degeneration caused by mutations in the *CHM* gene, which encodes Rab Escort Protein 1 (REP1), a protein thought to be involved in membrane trafficking.^19, 20^ Mutations in *CHM* lead to progressive degeneration of the photoreceptors, retinal pigment epithelium (RPE), and choroid.^21–24^ Patients with CHM typically present in their youth with nyctalopia and visual field defects. The earliest clinically detectable abnormalities include RPE demelanization, disruption of the photoreceptor outer segments, and severe rod photoreceptor dysfunction starting in the near mid-peripheral retina.^25–27^ Cone dysfunction as well as centrifugal and centripetal movement of the degenerative process^21, 27^ causes progressive constriction of the visual field and eventual involvement of the foveal center, which results in degraded visual acuity typically in the fifth decade of life.^21, 25, 27–29^

Retinal imaging in CHM has demonstrated retained central islands of neural retina, with sharp borders demarcating the transition to severely degenerated areas within or in the periphery of the retained islands.^29, 30^ Cross sectional imaging with optical coherence tomography (OCT) has revealed that, independent of the cellular origin of the primary mechanism of disease, shortening or loss of the photoreceptor outer segments are the earliest clinically detectable abnormalities, preceding overt structural loss within the RPE and choroid.^21, 24, 27, 29, 31^

At the cellular level, imaging with adaptive optics scanning laser ophthalmoscopy (AOSLO) has enabled visualization of the photoreceptor mosaic both in health and disease.^32^ This technique involves measuring and compensating for the optical aberrations of the living eye, in order to obtain diffraction limited imaging through the natural pupil of the eye.^33^ In CHM, AOSLO imaging has revealed the photoreceptor mosaic remains contiguous up to the edge of sharp borders between relatively preserved and atrophic retina.^29, 34, 35^ Within the retained central islands, local regions of the photoreceptor mosaic can exhibit either normal or reduced cone densities with dim and mottled waveguided reflectance profiles. AOSLO imaging combined with nearly cellular-level measures of vision, or adaptive optics (AO) microperimetry, has confirmed the existence of sharp transitions between functioning retina and severe sensitivity losses that collocate with the observed rapid transitions in retinal structure.^30^ The results suggest the RPE, in addition to rods, is an autonomous site of degeneration in CHM. Cones ultimately are lost as well, either by mechanisms that occur in parallel or as a consequence of severe RPE and/or choroidal changes.^21, 27, 29, 30, 34, 36–39^

Current gene augmentation strategies for treating CHM target these residual islands of viable retina hoping to maintain or improve levels of central vision that are often above the legal limit of blindness, albeit associated with very constricted visual fields.^21^ Delivering normal copies of the *CHM* gene aimed at restoring REP1 function is currently performed by a subretinal injection.^7, 17^ The procedure is not without risk, in that it involves forcefully delivering a fluid under the neural retina, thus separating the photoreceptors within the residual island from the underlying supportive RPE. This process of intentionally detaching the photoreceptors from the RPE has the potential to damage the structural integrity of the entire retina, particularly the RPE and photoreceptors. In the present study, we use AOSLO to gain insight into the short-term changes of the cone mosaic following macular subretinal injections of AAV2-*hCHM* in CHM subjects. We evaluate the cone mosaic structure in conjunction with foveal measures of cone outer segment length and vision prior to and one-month after the subfoveal injections.

## Methods

### Subjects and General Procedures

This research adhered to the Declaration of Helsinki and was approved by the Institutional Review Boards at the University of Pennsylvania and The Children’s Hospital of Philadelphia. All nine participants provided informed consent before voluntarily enrolling in the study. The subjects also provided informed consent and were enrolled in a dose escalation Phase 1/2 clinical trial testing the safety of the subretinal delivery of AAV2-*hCHM* in subjects with CHM (ClinicalTrials.gov identifier NCT02341807). Inclusion criteria included male gender, 18 years of age or older, confirmed disease-causing *CHM* gene mutation, central visual field constriction <30 degrees in at least one of 24 meridians, visual acuity better than 20/200, interocular symmetry in disease severity, exclusion of systemic or ocular diseases or medications that could potentially interfere with the disease process or delivery of the subretinal injection, and compliance with the clinical trial study protocol.

Subjects underwent a complete ophthalmic examination before and one month after the subretinal delivery of AAV2-*hCHM*, including dark-adapted foveal sensitivity testing, OCT and AOSLO imaging. Cone sensitivity was measured at the fovea using a modified Humprey Field Analyzer (HFA II-I, Carl Zeiss Meditec, Dublin, CA) using a 1.75 degree diameter, 650 nm stimuli presented at the foveal center following 30 minutes of dark adaptation.^21^ Axial lengths for both eyes were recorded using an IOL-Master® (Carl Zeiss Meditec, Dublin, CA). All AOSLO images were proportionally scaled by axial length as has been done previously. Within 87 days from the baseline imaging session (mean 27 ± 25 days, range 4 – 87 days), subjects then received unilateral subretinal injection of low dose (up to 5x10 vector genome (vg) per eye, n=5) or high dose (up to 1x10 vg per eye, n=4) AAV2-*hCHM* per the Phase 1/2 clinical trial protocol. As previously reported, the injection ‘blebs’ covered the entire extent of the residual central islands including the foveal center. The planned upper limit for the volume of the subretinal injections was 300 µl; the final volume injected was limited to that required to produce a visible subretinal bleb that covered the residual islands (between 20 µl and 100 µl).^16^ For quantitative analyses of the OCT cross-sections, images from post-operative visits were co-registered to their baseline, re-sampled at ten-fold the original resolution. Longitudinal reflectivity profiles (LRPs) from the foveal center (or juxtafovea in PN07 to avoid retinal tracks and EZ discontinuation) were generated using ImageJ imaging analysis software (http://imagej.nih.gov/ij/; provided in the public domain by the National Institutes of Health, Bethesda, MD, USA) following published methodology.^21, 40^ LRPs aligned by the main RPE/BrM signal peak were used to determine the inter-peak distance between the EZ signal peak to the peak at the base of the RPE/BrM. The distance corresponds to the combined length of the photoreceptor inner and outer segment as well as the height of the RPE or EZ-to-BrM distance. Finally, changes in dark-adapted cone sensitivity and foveal EZ-to-BrM distance were compared with potential changes in the cone mosaic morphology as determined by AOSLO imaging.

### AOSLO Imaging Procedures and Image Processing

The AOSLO system used in this study has been previously described.^41, 42^ Briefly, the custom-built, multi-modal AOSLO apparatus consisted of an 848 Δ26 nm superluminescent diode (SLD) for wavefront sensing and a 795 Δ15.3 nm SLD for near-infrared imaging (Superlum, Cork, Ireland). Wavefront correction was performed using a 97-channel deformable mirror (Alpao SAS, France). Three photomultiplier tubes (Hamamatsu Corporation) were configured to record confocal and non-confocal split-detection image sequences at 18 Hz simultaneously. CHM patients were aligned to the custom-built AOSLO imaging system using a dental impression. Patients were instructed to fixate at a target using the imaging eye. AOSLO image sequences were acquired using both a 1.75° and 1° square imaging field over the central 3 degrees surrounding fixation and using a 1 square° imaging field from fixation out along each meridian until reaching the atrophic lesion border or reaching ∼15° eccentricity. The custom-built AOSLO allowed higher resolution imaging (theoretical limit ∼2 µm) for visualization of waveguiding foveal and parafoveal cones as well as the split-detection modality for visualization of the cone inner segments.

Image sequences from the custom built AOSLO were desinusioded, and several reference frames were chosen automatically from each image sequence using a custom MATLAB algorithm based on the method published by Salmon et al.^43^ Custom software was used to remove intra-frame distortions caused by eye motion and 50 frames of the AOSLO image sequence were registered.^44^ Registered frames were then averaged together and the averaged image was dedistorted using a custom MATLAB algorithm based on Bedggood and Metha^45^ to remove distortions caused by eye motion within the reference frame. The dedistorted images acquired from within a single imaging session were then automatically montaged using MATLAB as previously described.^46^ This montaging step of the analysis was supplemented with manual image alignments at retinal locations where an automated match was not found, but an image was acquired. AOSLO image montages from each time point were then manually aligned to each other longitudinally using Adobe Photoshop. Macroscopic image features, such as blood vessels and the contours of the central island, from the full montage were used for a gross, initial longitudinal alignment; the longitudinal alignment was then refined for cone-by-cone accuracy over regions of interest (ROIs). When necessary, one-month images were scaled to the baseline images.

### Cone Density Measurements

Four ROIs from each eye were selected for measurement of cone densities. ROIs were manually cropped from both the baseline and one-month post-injection split detection AOSLO montages for both injected and control eyes of all subjects. Cones were manually identified using custom software. One grader, JIWM, identified cones in all ROIs. The grader was masked to treated vs control eye and time point for each subject. The grader was able to adjust the brightness and contrast of the image both in linear and logarithmic displays while selecting cones within the ROIs. Cone centers were used to determine the Voronoi boundaries for each selected cone and bound cone density was calculated for each ROI.^47^ Cone densities were compared between control and injected eyes at each time point and between time points for control and injected eyes. Paired t-tests were used to determine statistical differences at p<0.05.

## Results

Nine molecularly confirmed CHM subjects participated in the study. Subject characteristics are shown in **Table 1**. Subjects ranged in age from 26-50 years at the time of enrollment. As previously reported, surgeries were uneventful.^16^ Axial lengths ranged from 23.33 – 26.95 mm (mean ± standard deviation: 25.01 ± 1.21 mm). Foveal cone sensitivity was unchanged at one-month post injection for 8 of 9 injected eyes and all 9 control eyes. One subject, PN-11 showed a significant loss in foveal cone sensitivity in the injected eye (**Table 1**). As previously reported,^16^ this same subject demonstrated a significant loss in visual acuity (3 lines) in the injected eye while all other visual acuities remained unchanged.

**Table 1:** Study Subject Characteristics

|  | Study ID | Study ID* | Age (years) | Visual Acuity<sup>†</sup> |  | Injected Eye | Volume (μl)<sup>†</sup> | Axial length (mm) |  | *CHM* Mutation | Change in Cone Sensitivity (dB) |  |
| --- | --- | --- | --- | --- | --- | --- | --- | --- | --- | --- | --- | --- |
|  |  |  |  | OD | OS |  |  | OD | OS |  | OD | OS |
| Low Dose Group<br>5x10<sup>10</sup> vg | PN-01 | 13057 | 33 | 20/40 | 20/32 | OD | 50 | 26.81 | 26.95 | c.745del.T | -1 | -2 |
|  | PN-03 | 13125 | 32 | 20/32 | 20/25 | OD | 50 | 23.44 | 23.33 | c.1437dupA | 1 | 4 |
|  | PN-04 | 13071 | 33 | 20/25 | 20/25 | OS | 120 | 23.83 | 23.97 | c.1663A>T | 5 | 0 |
|  | PN-05 | 13004 | 50 | 20/40 | 20/20 | OD | 20 | 26.56 | 26.85 | Large Exon 1 deletion | -4 | 6 |
|  | PN-06 | 13131 | 37 | 20/25 | 20/25 | OS | 25 | 24.99 | 24.76 | c.1327_1328delAT | -2 | -2 |
| High Dose Group<br>1x10<sup>11</sup> vg | PN-07 | 13159 | 43 | 20/25 | 20/20 | OD | 100 | 25.39 | 25.25 | c.1144G>T | 2 | 2 |
|  | PN-08 | 13039 | 26 | 20/25 | 20/25 | OD | 50 | 25.78 | 25.76 | c.1327_1328delAT | 0 | 1 |
|  | PN-09 | 13224 | 57 | 20/20 | 20/20 | OD | 100 | 24.30 | 24.36 | c.41dupT | -12 | 3 |
|  | PN-11 | 13226 | 39 | 20/20 | 20/20 | OD | 100 | 24.06 | 23.85 | c.940-2A>T | -3 | -1 |

\* ID used in previous cross-sectional studies conducted by our research group.<sup>21,29,30,72</sup>
<sup>†</sup> Previously reported by Aleman et al.<sup>16</sup>

### Shortening of Cone Outer Segments after Subretinal Gene Therapy

Overall, at one-month post-injection OCTs show that compared to baseline images the laminar architecture of the retina is qualitatively unchanged in both the injected and uninjected eyes and that the subretinal bleb containing AAV2-*hCHM* has resolved in injected eyes (**Figure 1**). Quantitative analyses showed a normal foveal ellipsoid zone (EZ)-to-RPE/Bruch’s membrane (BrM) distance (mean normal +2SD = 52 ± 15 µm) at baseline in all subjects, except PN05 and PN07 with the most severe foveal abnormalities. At the one-month post-injection time point, however, there was shortening of this distance in the AAV2-*hCHM* injected eyes, suggestive of foveal cone outer segment shortening (**Figure 1C**). The differences between the measures at one-month and baseline exceeded the variability of the measurements (± 4.42 µm) (Figure 1C, dashed lines) in four subjects, was borderline significant in another four subjects, and unchanged in one subject. The EZ-to-RPE distance in uninjected eyes remained unchanged at the one-month time point compared to baseline measurements.

**Figure 1:**
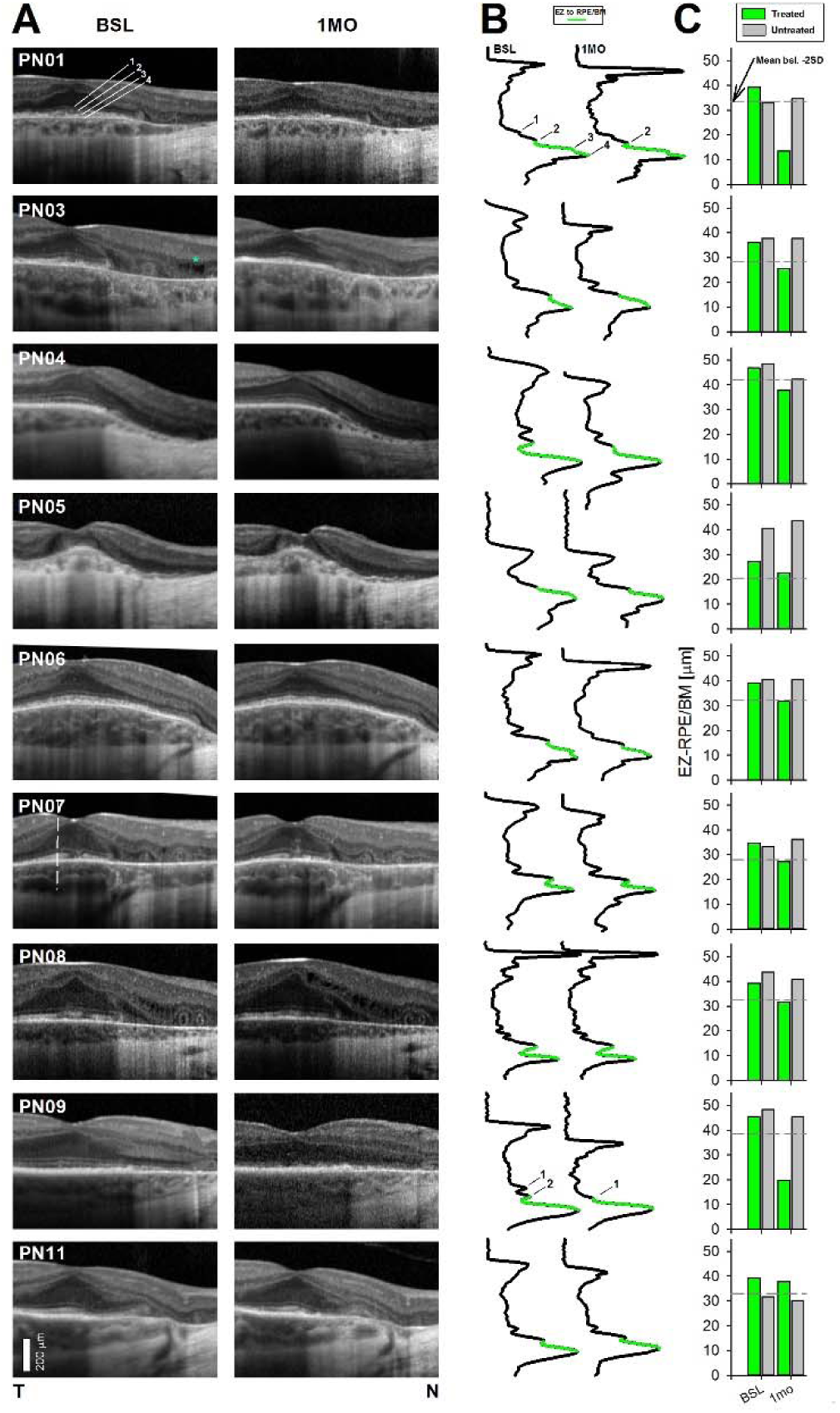
Acute outer retinal changes after the subfoveal subretinal injections. 3 mm SD-OCT horizontal SD-OCT cross-sections through the fovea at baseline compared to 1 month after the subretinal injections in the injected eyes (**A**). Outer photoreceptor laminae are labeled in PN01 following convention: 1. external limiting membrane (ELM), 2. Inner segment ellipsoid zone band (EZ); 3. interdigitation (IZ) of the outer segment with the apical RPE; 4. retinal pigment epithelium/Bruch’s membrane (RPE/BrM) band. Longitudinal reflectivity profiles (LRPs) from the fovea at 1 month post-injections compared to baseline (**B**). Segment colored green on the LRPs is the distance from the EZ to RPE/BrM, which relates to the photoreceptor outer segment (POS) length. EZ-to-RPE/BrM distance in the study eye compared to the control eye for each subject (**C**). Dashed lines define mean-2SD of the intersession variability of the measures in uninjected CHM eyes as well as from estimates in normal subjects, which defines limit for significant thinning of the injected eye.^40^ Green bars represent injected eyes, gray bars are uninjected eyes. The EZ-to-RPE distance was decreased at one-month post-injection compared to baseline measurements; this shortening was significant in four subjects (PN-01, PN-03, PN-04, PN-09), borderline significant in another four (PN-05, PN-06, PN-07, PN-08), and unchanged in one subject (PN-11).

### Photoreceptor Mosaic Integrity and Cone Photoreceptor Density

Non-confocal split detection AOSLO at baseline revealed the photoreceptor mosaic within the central island of remaining retina was intact and contiguous out to the border of atrophy, at which point, the photoreceptor mosaic exhibited a sharp transition to atrophic retinal regions. AOSLO at one-month post-injection also revealed a contiguous mosaic in 8 of 9 injected eyes and all 9 uninjected eyes (**Figure 2, Supplemental Figures 1**-**8**). Global features within the AOSLO montage and photoreceptor mosaic could be aligned longitudinally between time points. Local distortions in adjacent images both within and between timepoints however, precluded cone-by-cone alignment across the full montage in both injected and uninjected eyes. Thus, ROIs within the cone mosaic were selected for cone-by-cone alignment across time points using rigid transforms (translation, rotation, scale) only (**Supplemental Figure 2**). Cone-by-cone alignment was attained at multiple retinal locations within the montage in all eyes, including both injected and uninjected eyes. Qualitatively, this manual alignment was easier to perform in uninjected eyes and ROIs in uninjected eyes showed accurate cellular alignments over a larger distance than ROIs in injected eyes.

**Figure 2:**
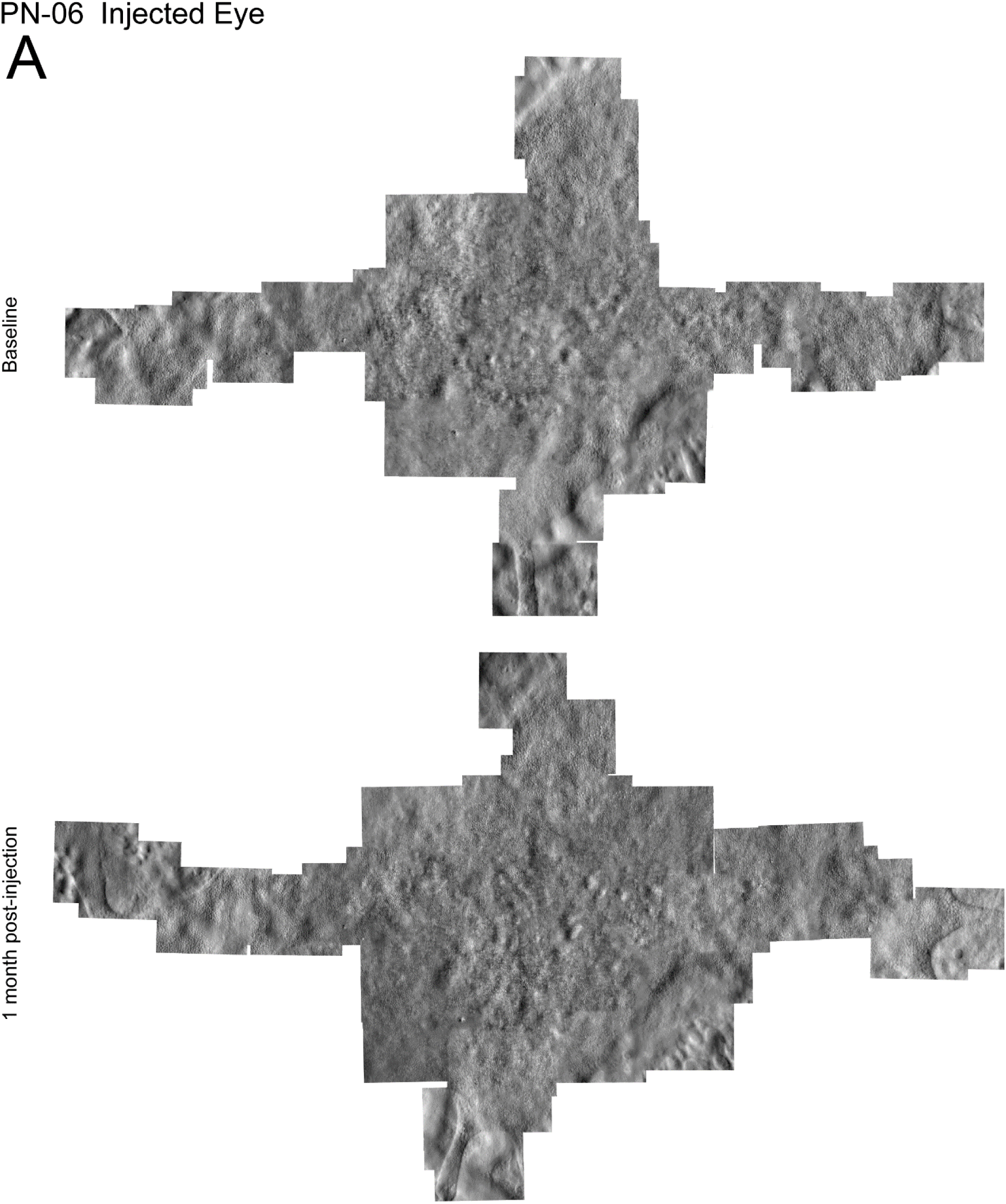
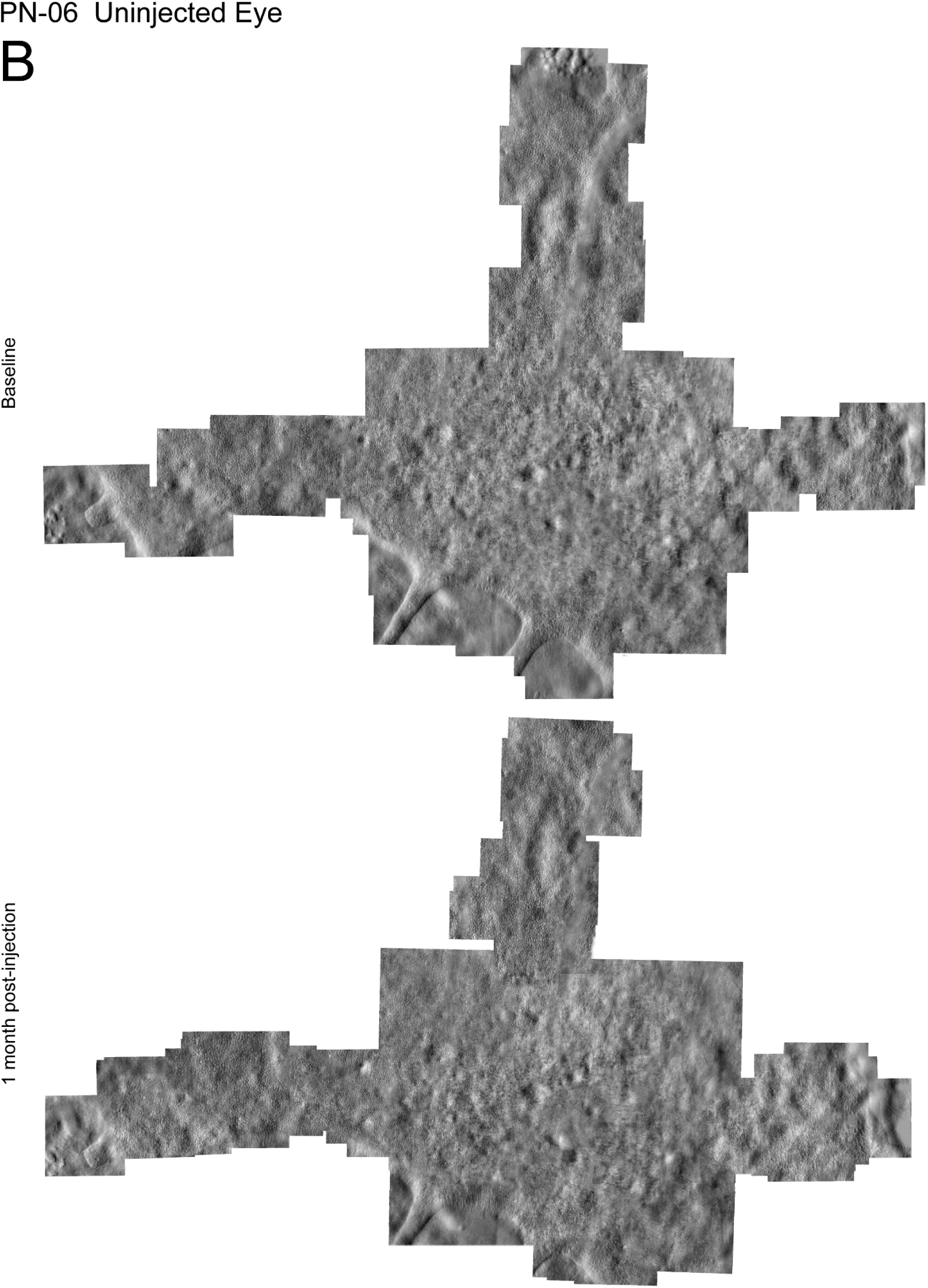
Nonconfocal split detection AOSLO montage of the photoreceptor inner segment mosaic at baseline and one-month post-injection in the injected (**A**) and uninjected (**B**) eyes of PN-06. The same retinal features are observed longitudinally in both eyes and the photoreceptor mosaic remains intact following the subretinal injection of AAV2.*hCHM*.

Cones could be manually identified and bound cone density determined for longitudinally aligned ROIs in all eyes (**Figure 3**). Cone densities were similar across timepoints for all ROIs in both injected and uninjected eyes (**Figure 4**). In injected eyes, cone density (mean ± standard error) was 24,027 ± 1,991 cones/mm^2^ at one-month post-injection compared to 24,401 ± 2,361 cones/mm^2^ at baseline. Cone densities in uninjected eyes were 24,284 ± 3,051 cones/mm^2^ at one-month compared to 24,491 ± 3,022 cones/mm^2^ at baseline (**Table 2**). Summarizing across all ROIs, there was no statistical difference observed between cone densities measured in injected and uninjected eyes at either timepoint (P= 0.97, 0.91 for baseline and one-month time points respectively). There was no significant difference in the difference in cone density between injected and uninjected eyes (P=0.80) and no significant difference in the percent differences in cone density between injected and uninjected eyes (P=0.60).

**Figure 3:**
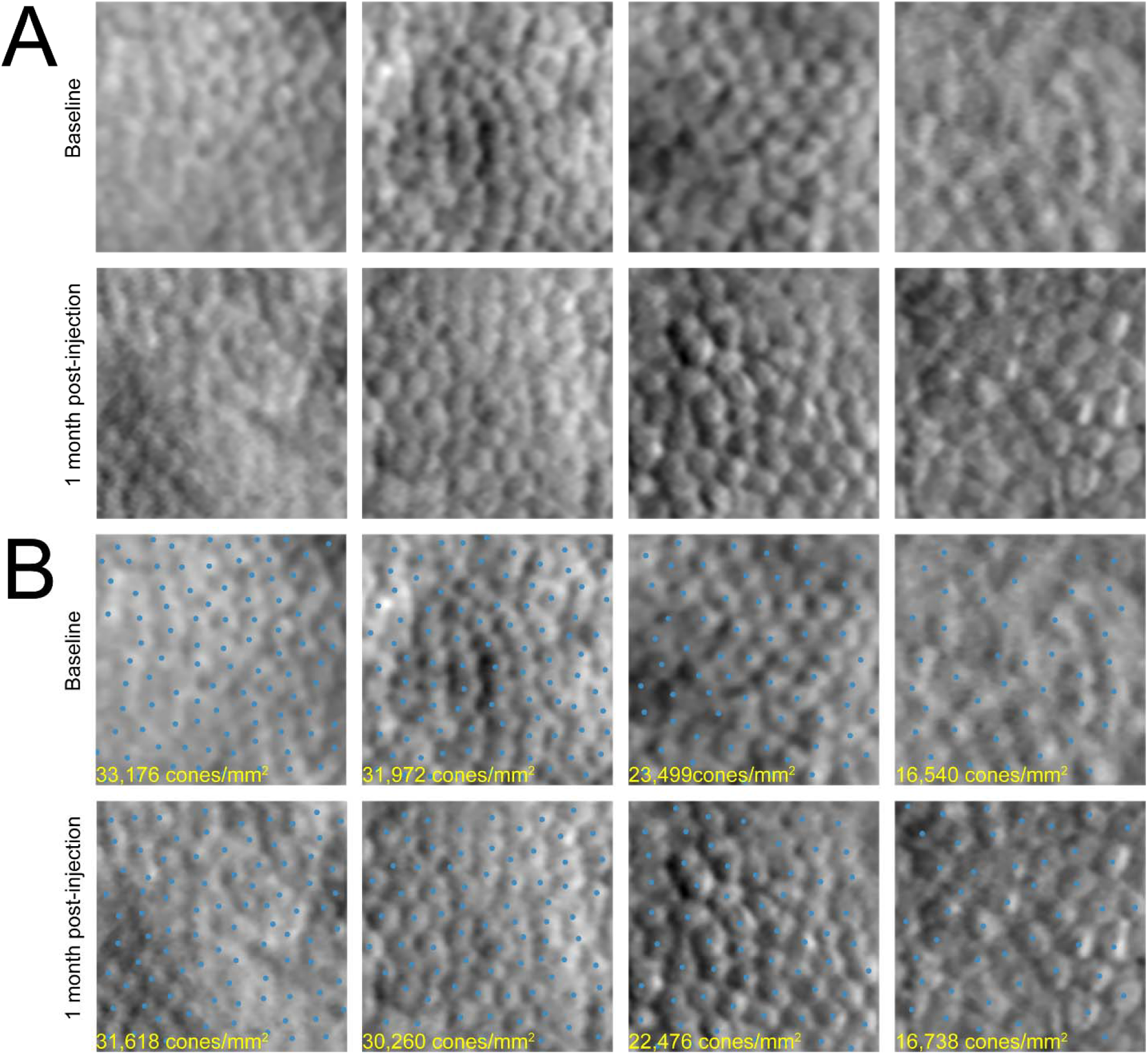
A: Adaptive optics (AO) regions of interest (ROIs) aligned between time points (top baseline, bottom one-month post-injection) from the injected eye (OS) of subject PN-06. An intact cone mosaic is visible before and after the subretinal injection of AAV2-*hCHM*. **B:** Cones were manually identified (blue dots) and bound cone density was calculated for each ROI (cone densities in yellow for each ROI). No significant changes in cone density were observed between baseline and one month measurements.

**Figure 4:**
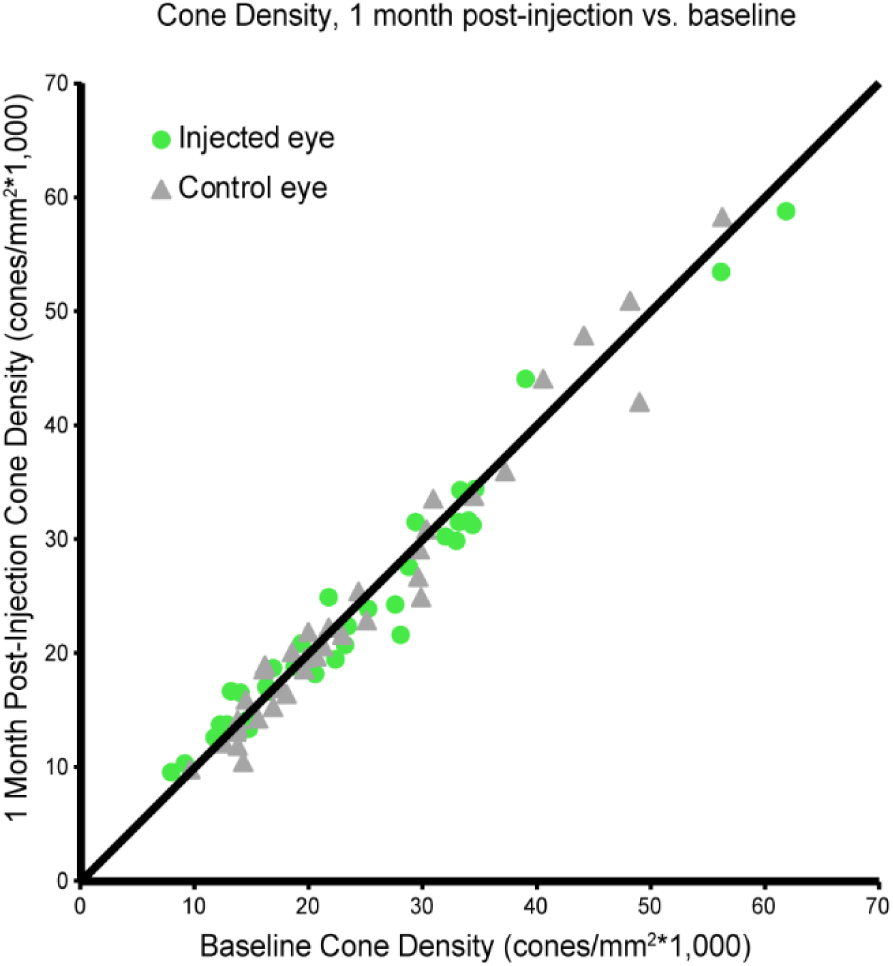
Cone density at one-month post-injection versus baseline. There was no statistical difference between one-month post-injection measurements and baseline measurements in either injected or control eyes. Green circles: injected eyes, Gray triangles: control eyes. Black line is the line of equivalence.

**Table 2:**
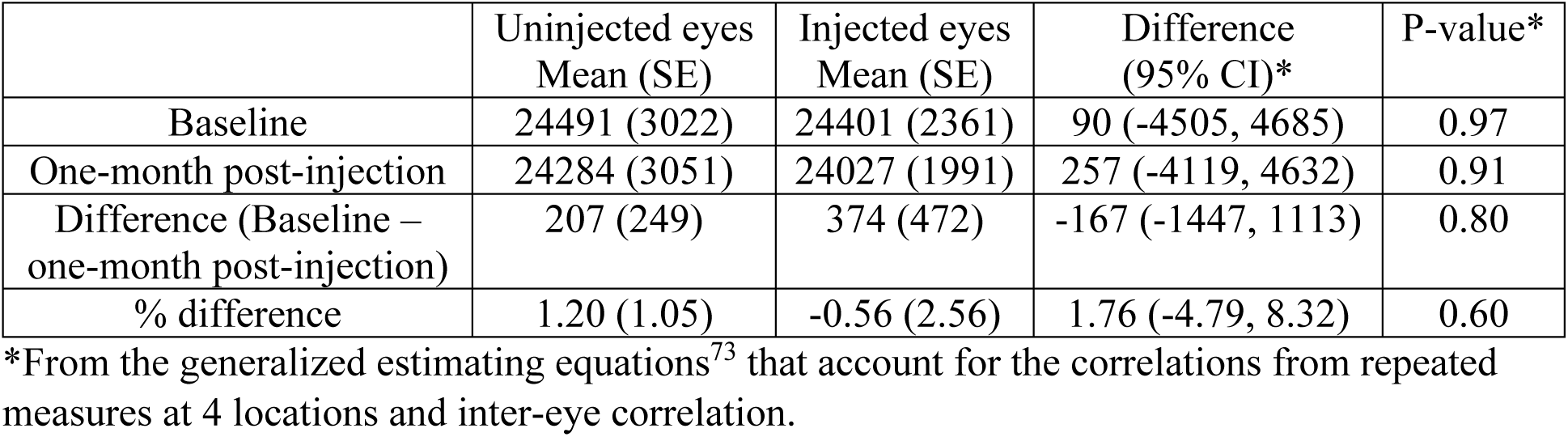
Comparison of cone density (cones/mm^2^) between injected and uninjected eyes (n=9 subjects, 18 eyes)

One-month post-injection AO images of subject PN-09 revealed a local loss of cones at the fovea (**Figure 5**). This area of cone loss was co-located with the retinal region that revealed a loss of foveal sensitivity at the one-month post-injection timepoint. Despite the foveal cone loss, cones were visible in the surrounding parafoveal regions at the same time point. Similar to the other subjects, the photoreceptor mosaic was intact at regions outside of the fovea for PN-09, longitudinal alignments of the cone mosaic were possible, and cone densities were unchanged in ROIs selected for cone density quantifications.

**Figure 5:**
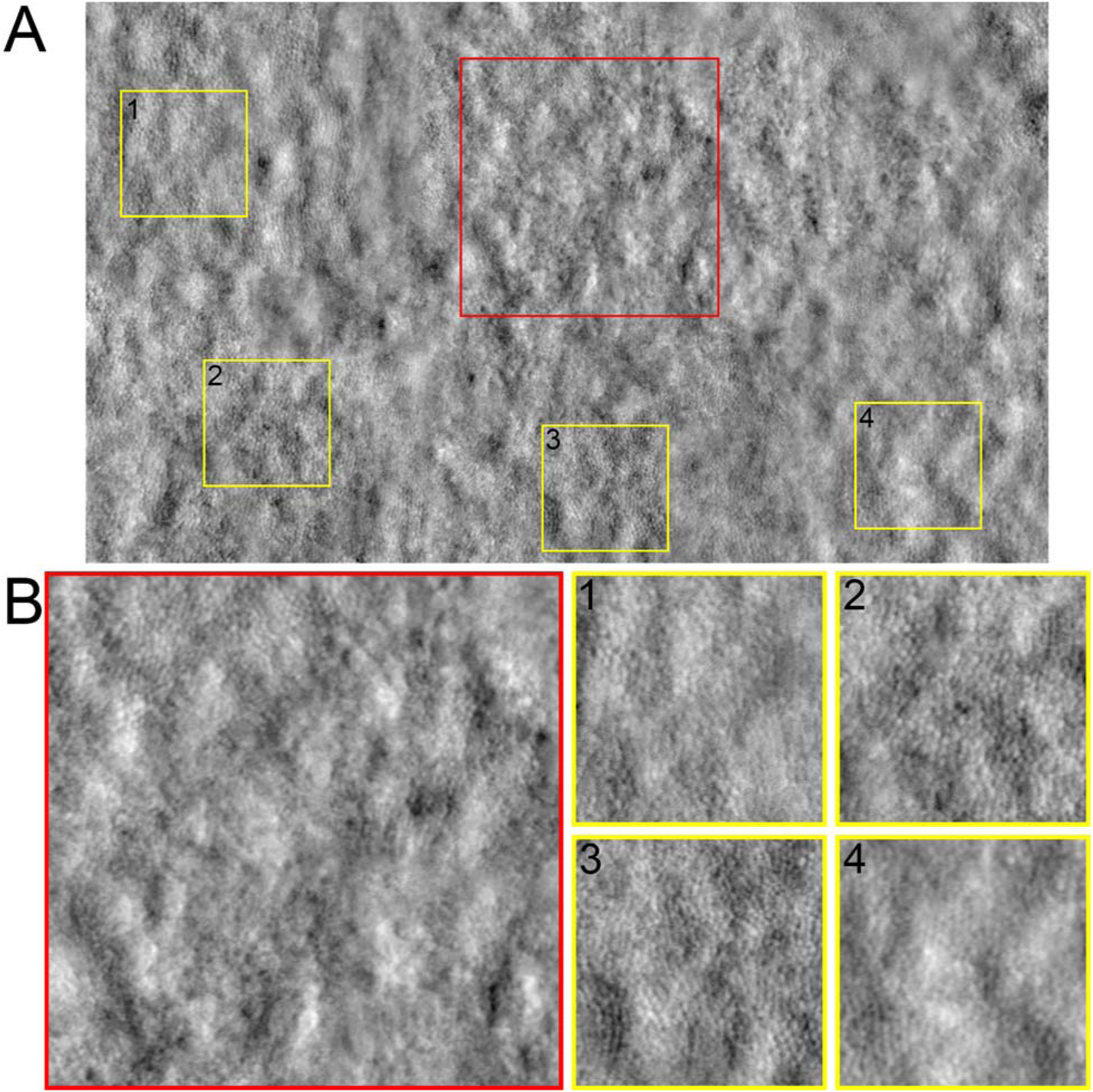
AO nonconfocal split detection montage (**A**) of the foveal region in the injected eye of PN-09 at one-month post-injection. **B**: Enlarged (2x) regions of interest showing a loss of cone inner segments at the fovea (red box) surrounded by an intact cone inner segment mosaic in the parafoveal regions (yellow boxes).

## Discussion

Treating CHM by gene augmentation represents a significant departure from the treatment scenario of the earlier experience in *RPE65*-IRDs that culminated with the first FDA-approved gene therapy product for use in the clinic.^3, 18^ In mid- to end-stage CHM, there is no alternative but to treat small fragile central islands of relatively preserved retina that sustain limited fields of vision that often support reading levels that are above the legal limit of blindness. The scenario is quite different from the treatment of functionally blind retinas that are less fragile in diseases within the spectrum of *RPE65*-LCA.^3, 18^ In fact, the resulting shift in the benefit-to-risk ratio is to be expected for the larger group of non-LCA IRDs at the disease stages that are typically considered for initial clinical trials, making CHM a model for an entire group of genetic retinal diseases that await treatment solutions. Although the safety profile of the subretinal injections is now better understood, quantitative approaches, particularly at the cellular level, are needed to understand the mechanisms that lead to unwanted outcomes, particularly when the remaining vision is threatened by an invasive procedure, such as a therapeutic retinal detachment. In the current study we used OCT and AOSLO imaging to document possible short-term changes of the photoreceptor mosaic following subfoveal injections of AAV2-*hCHM*. We demonstrate that at one-month post-injection the cone mosaic resettled on the RPE following resolution of the subretinal bleb. The cone mosaic remained intact in eight of nine subjects (except PN09), and outside of the foveal center in all subjects. Quantification of cone densities revealed no measurable difference between the injected and uninjected eyes and no measurable changes in cone densities between baseline and one-month post-injection. These results show there was no significant widespread cone loss across the retained area of central retina targeted by the retinal detachment in any of the nine CHM subjects included in this study. Thus, we make two important conclusions regarding the safety of the AAV2-*hCHM* experimental therapy. First, cone photoreceptors did not drop out as a consequence of mechanical or acute inflammatory changes in response to the presence of AAV2-*hCHM* in the subretinal space. Second, the therapeutic retinal detachment, as performed by us, did not result in detectable short-term changes of the density of the photoreceptor mosaic, although mild shortening of the photoreceptor outer segments was detected. Altogether, our results provide safety information at the cellular level for both the surgical technique and the AAV2-*hCHM* study agent confirming histologic/cellular-level safety signals that up to now were only available through histopathologic studies in normal non-human primates.^48–52^

The subretinal injection however, is not without risk. Foveal thinning and the occurrence of full thickness loss of retinal tissue (macular holes) are known complications of the procedure.^3^ The loss of the photoreceptors at the fovea in one of our study subjects (PN09) raises the possibility of individual vulnerability to the subfoveal injection, an issue reported in at least one subject in each of the CHM gene therapy clinical trial reports.^18^ PN-09’s surgery was considered uneventful. It is unclear why the photoreceptors at the fovea of PN-09 did not withstand the subfoveal injection, however, because the parafoveal cones did survive the intervention, we suggest that the cone loss may have resulted from mechanical factors of the surgery rather than toxicity to the study agent. All study subjects were at a similar stage of their central retinal disease with remodeled foveas that were within normal limits of thickness (except PN05) or even thicker than normal.^21, 27^ In fact, two patients (PN07 and PN08) showed proximity of the transitional zones of structural disorganization to the foveal center, a factor known to predict the decline in visual acuity as part of the natural history of the disease,^21^ yet they did not have an unfavorable outcome.^16^ Inspection of PN09’s OCT cross-sections reveals faint EZ and IZ signal that may be indicative of fragile or more abnormal photoreceptor inner and outer segments, as has been described in certain forms of cone photoreceptor inherited degenerations, or cone and cone-rod dystrophies.^53, 54^ Perhaps these may be structural signs that dictate modified surgical approaches. Efforts to de-risk the subretinal injections with the introduction of mechanical devices that deliver precise volumes at prescribed hydrostatic pressures, as well as with the use of intraoperative SD-OCT systems that allow real time view of the microscopic retinal detail during the surgical interventions are some of the options.^55, 56^ Further studies are needed to address predisposing factors in patients with similar outcomes, as well as to determine the impact that alternative approaches to deliver gene therapy products to the desired cellular targets have on the health of the foveal photoreceptors.

Our OCT results at one-month post-injection showed a decrease in outer segment length in comparison to baseline thickness measurements. Preliminary results reported from Clinical Trial NCT02341807 showed that the foveal thickness slowly recovers over 6 months post-injection.^16^ This, taken together with the AO imaging results, leads to the conclusion that the measured decrease in thickness is caused by short-term outer segment shortening as opposed to cone loss. We hypothesize the forced detachment between the outer segment tips and the RPE from the surgical injection rather than the vector itself causes this short-term shortening of the outer segments. For unknown reasons the recovery of the outer segment length after retinal detachments follows a time course that defers from the normal renewal rate of the outer segment, and seems to be independent of the cause of the retinal detachment, whether short-lived and intentional, such as during the delivery of gene therapy products by subretinal injections, or after spontaneous primary macula-off detachments.^57–59^ Beyond purely mechanical factors affecting the outer retina, complex interactions are known to occur after retinal detachments, which may include the modulation of the structural outcome by variables inherent to the preexisting retinal degeneration, such as the response of the inner retina and the underlying RPE to the therapeutic detachment, topics in need of investigation.^60–63^

Although there have been reports documenting the safety of subretinal injections targeting the macula in CHM and other IRDs, this work is, to our knowledge, the first study to apply AO ophthalmoscopy to investigate the effect of an experimental gene therapy intervention for a blinding disease. Gene therapy aims to prevent cellular death and/or restore function to cells that are still surviving the genetic abnormality and AO ophthalmoscopy enables noninvasive visualization of individual cells. As a practical matter, the design and economics of gene therapy clinical trials puts a high value on accurately measuring outcomes reasonably soon after the experimental interventions. The Spark-funded clinical trial for CHM did not include AO imaging as an outcome measure, but the results from this ancillary study suggest that AO might be suitable as a precise anatomic outcome measure in future trials involving subretinal injections. Indeed, AO ophthalmoscopy is uniquely positioned to answer this need by allowing in-vivo and non-invasive visualization of the cellular targets of such interventions, particularly in disease states, such as neurodegenerations, where biopsies or other biologic markers of treatment effects are not available. Further, techniques such as optoretinography^64–68^ and AO microperimetry^30, 69–71^ have complemented AO imaging by allowing the direct or indirect evaluation of photoreceptor visual function at the cellular level. The emergence of these tools may prove impactful for assessing the short- and long-term safety and efficacy of gene therapies for blinding diseases.

In conclusion, our data support the short-term safety of subretinal injections of AAV2-*hCHM* as a treatment of CHM. Additional follow up will be required to assess the long-term safety and efficacy of subretinal injection and delivery of AAV2-*hCHM* for preventing or restoring vision loss caused by CHM.

## Acknowledgements

We thank Alfredo Dubra for sharing the adaptive optics scanning laser ophthalmoscopy optical design, as well as adaptive optics control, image acquisition and image registration software. We thank Gui-shuang Ying for help with statistical tests. **Funding provided by:** National Institutes of Health (NIH R01EY028601, R01EY030227, P30EY001583), Foundation Fighting Blindness, F. M. Kirby Foundation, Research to Prevent Blindness, the Center for Advanced Retinal & Ocular Therapeutics, and the Paul and Evanina Mackall Foundation Trust. **Competing interests:** AMM receives funding from Spark Therapeutics and REGENXBIO. JB also received clinical trial support from Spark Therapeutics. JIWM is an inventor on US Patent 8226236, US Patent App 16/389,942 and receives funding from AGTC. All data is available by request to the corresponding author.

## Supplemental material

Supplemental Figure 1-8: Nonconfocal split detection AOSLO montage of the photoreceptor inner segment mosaic at baseline and one-month post-injection in the injected (**A**) and uninjected (**B**) eyes of PN-01 (**Supplemental Figure 1**), PN-03 (**Supplemental Figure 2**), PN-04 (**Supplemental Figure 3**), PN-05 (**Supplemental Figure 4**), PN-07 (**Supplemental Figure 5**), PN-08 (**Supplemental Figure 6**), PN-09 (**Supplemental Figure 7**), PN-11 (**Supplemental Figure 8**).

**Supplemental Figure 1A.**
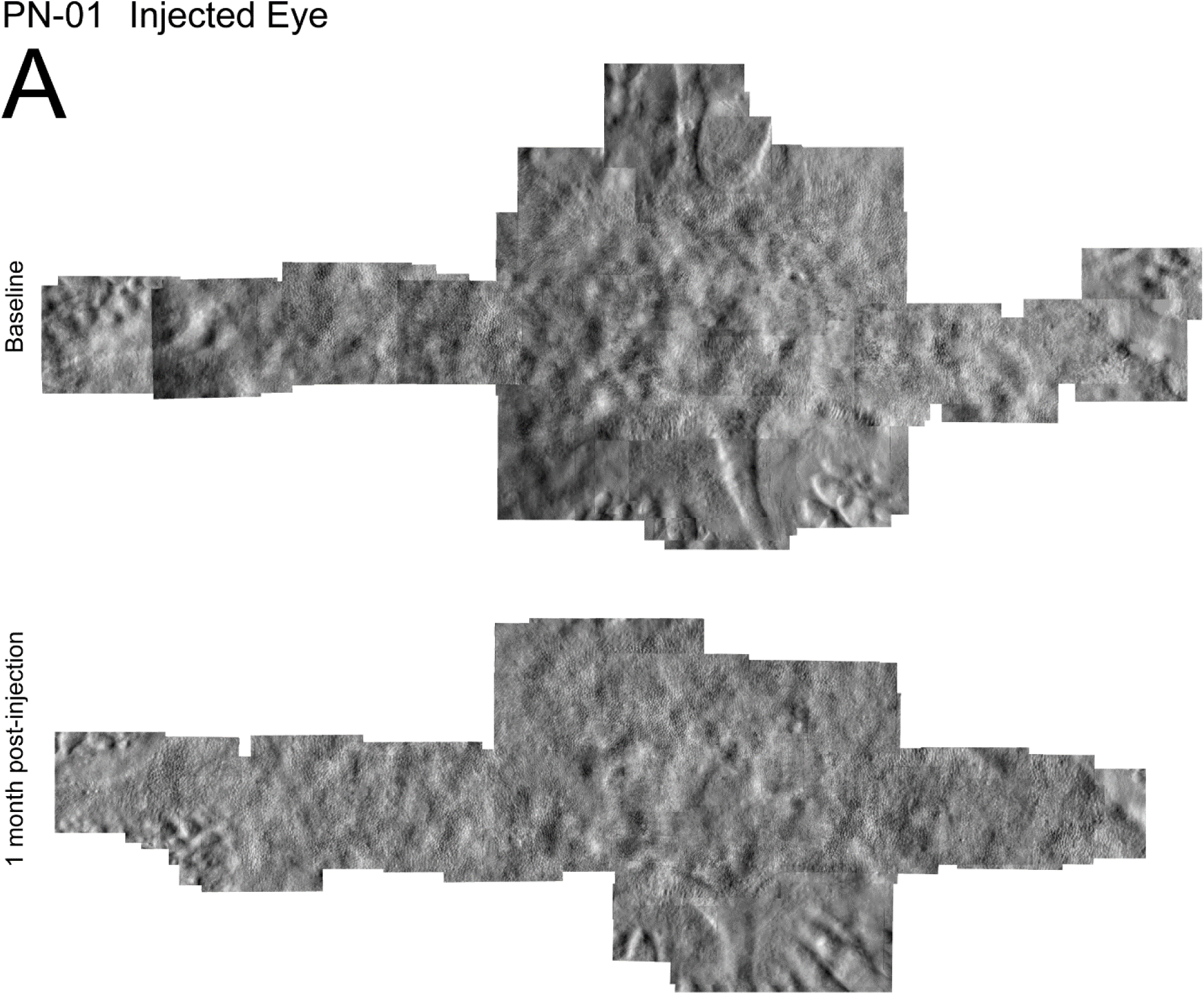

**Supplemental Figure 1B.**
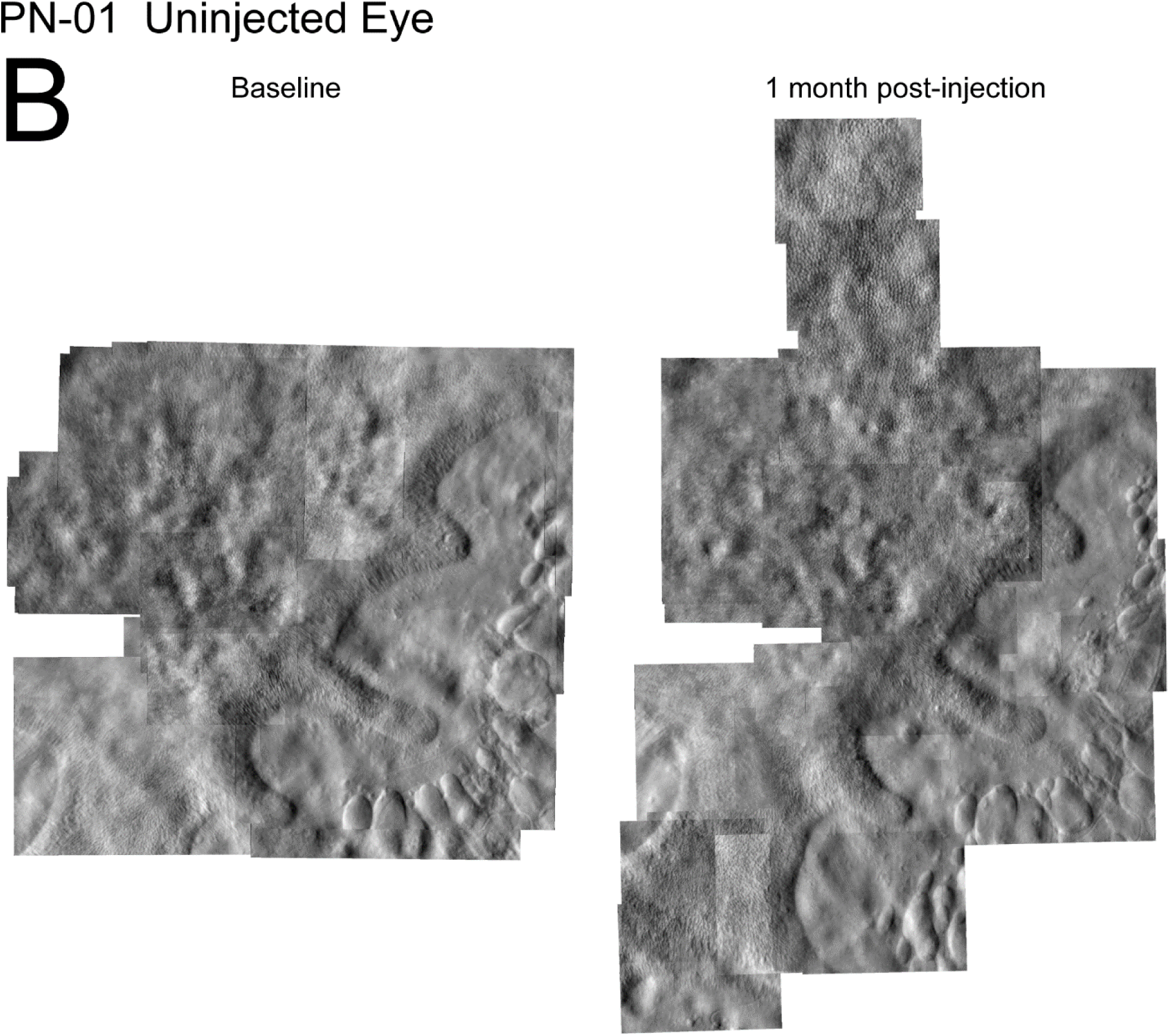

**Supplemental Figure 2A.**
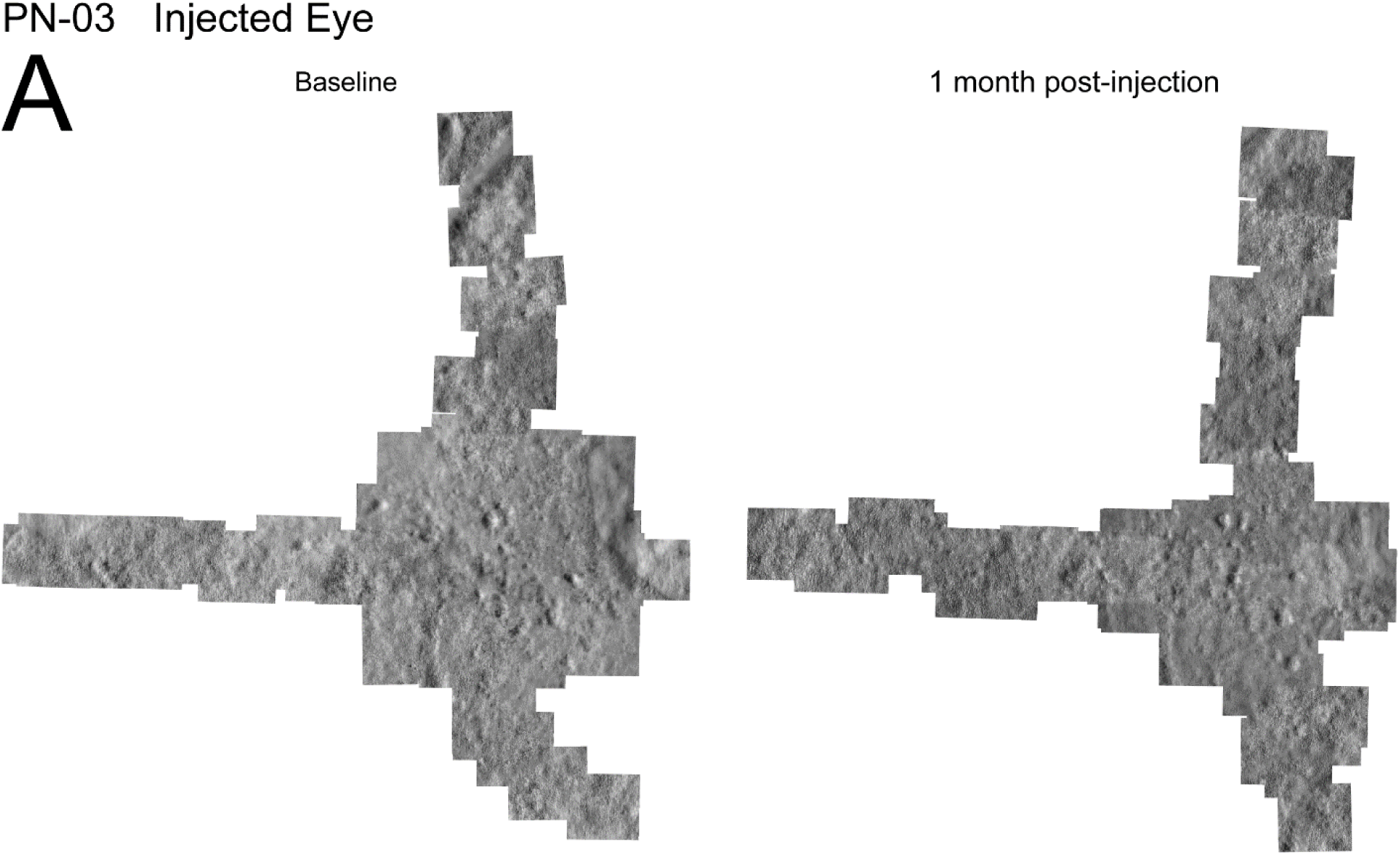

**Supplemental Figure 2B.**
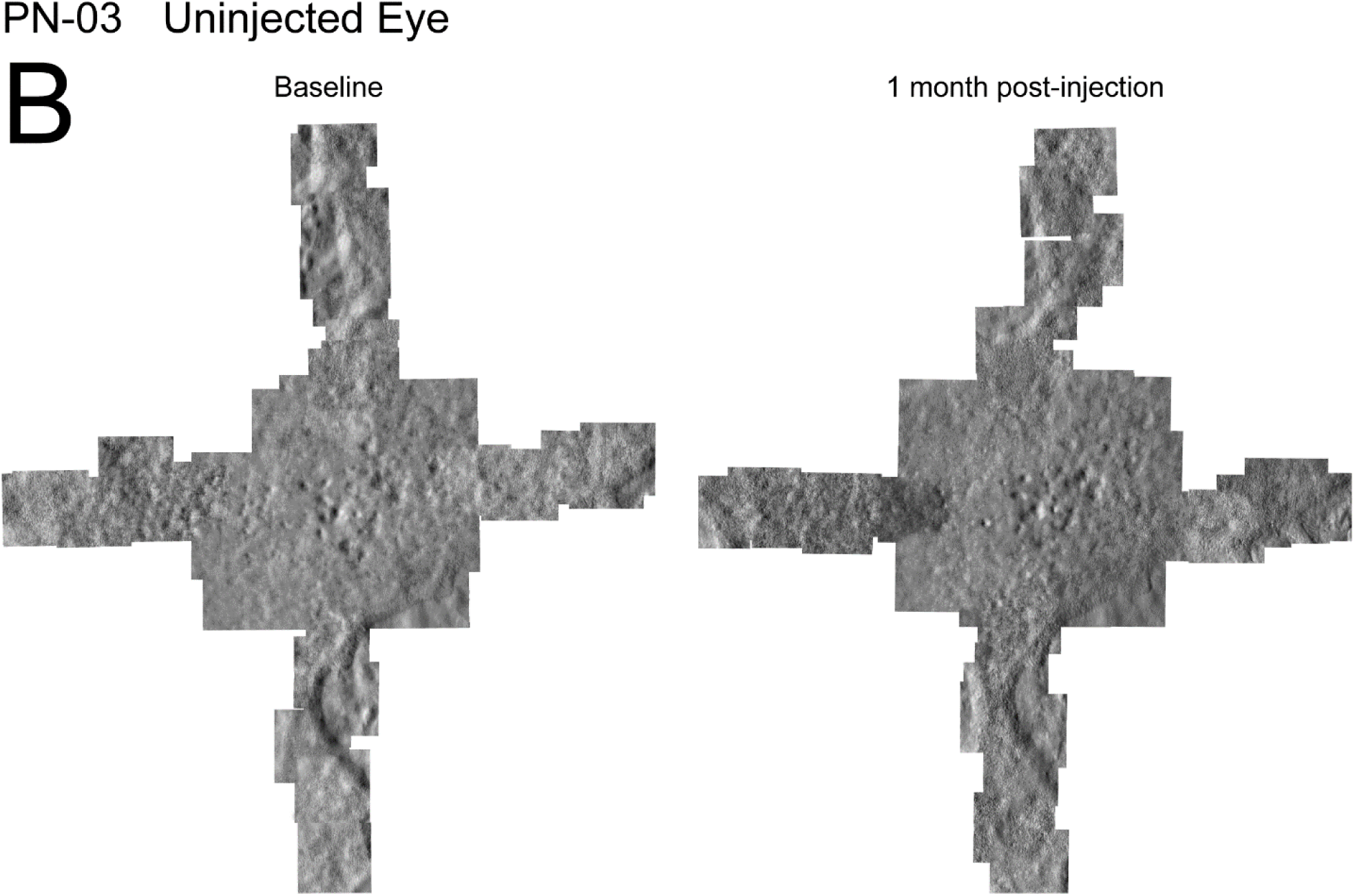

**Supplemental Figure 3A.**
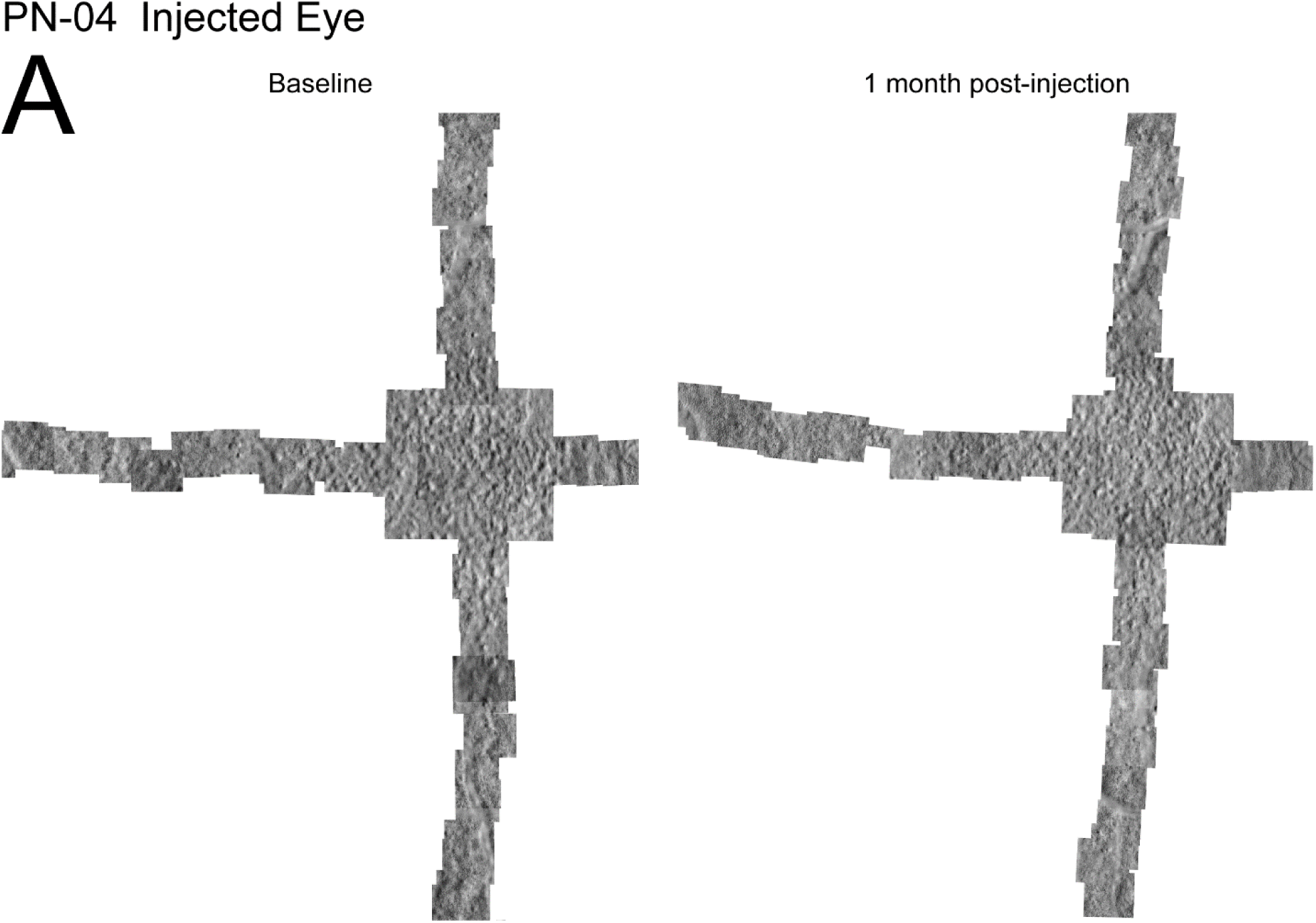

**Supplemental Figure 3B.**
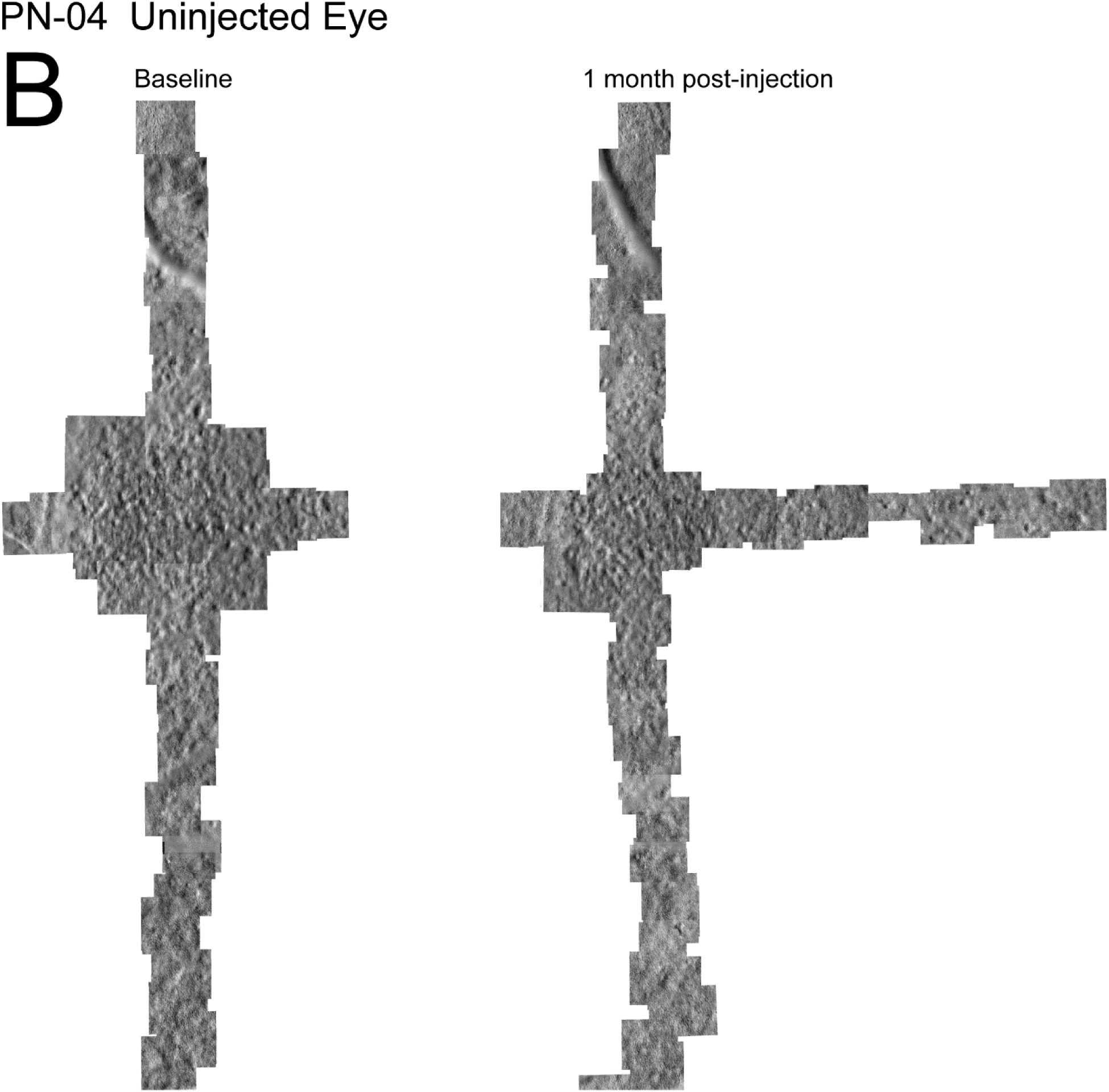

**Supplemental Figure 4A.**
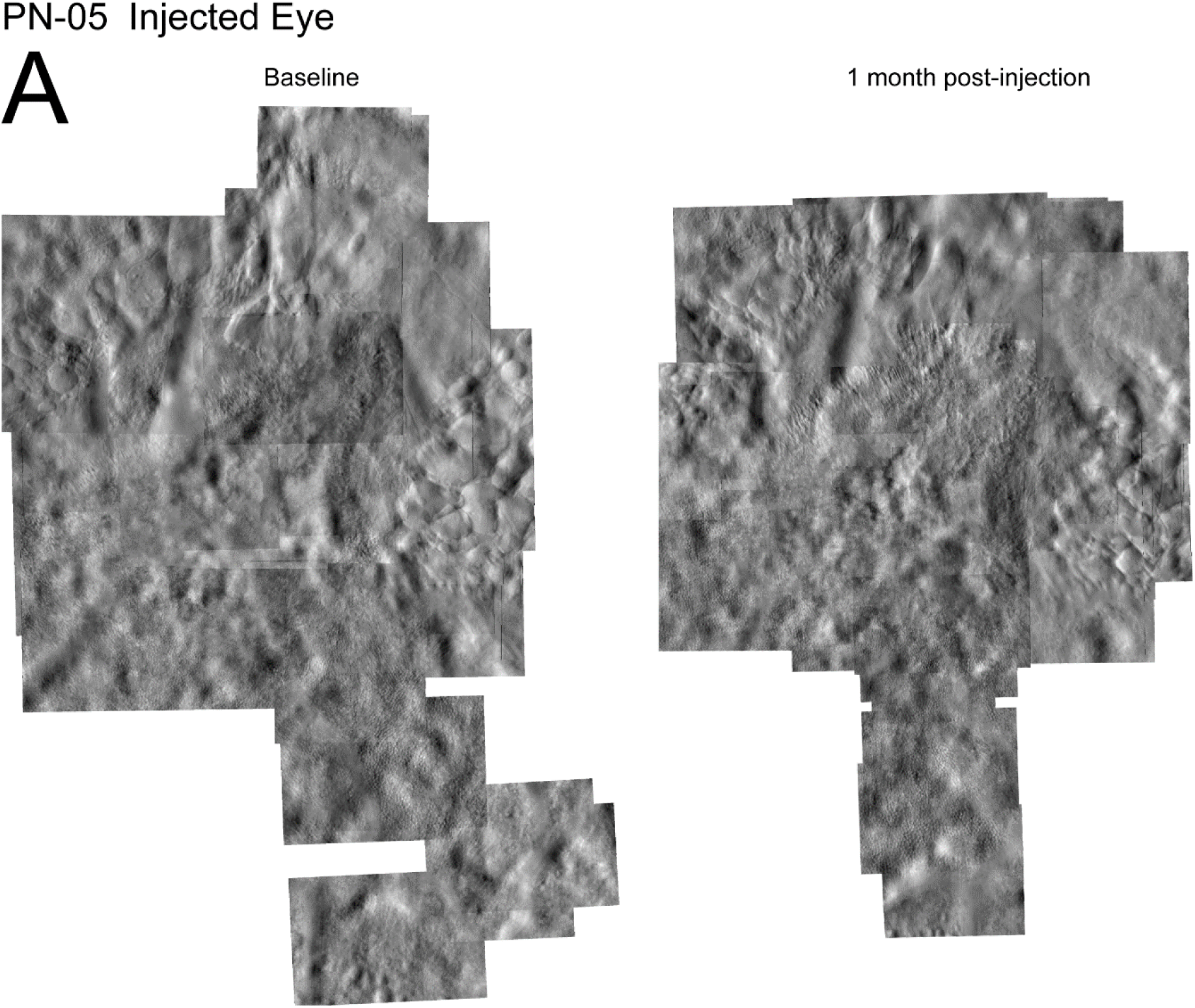

**Supplemental Figure 4B.**
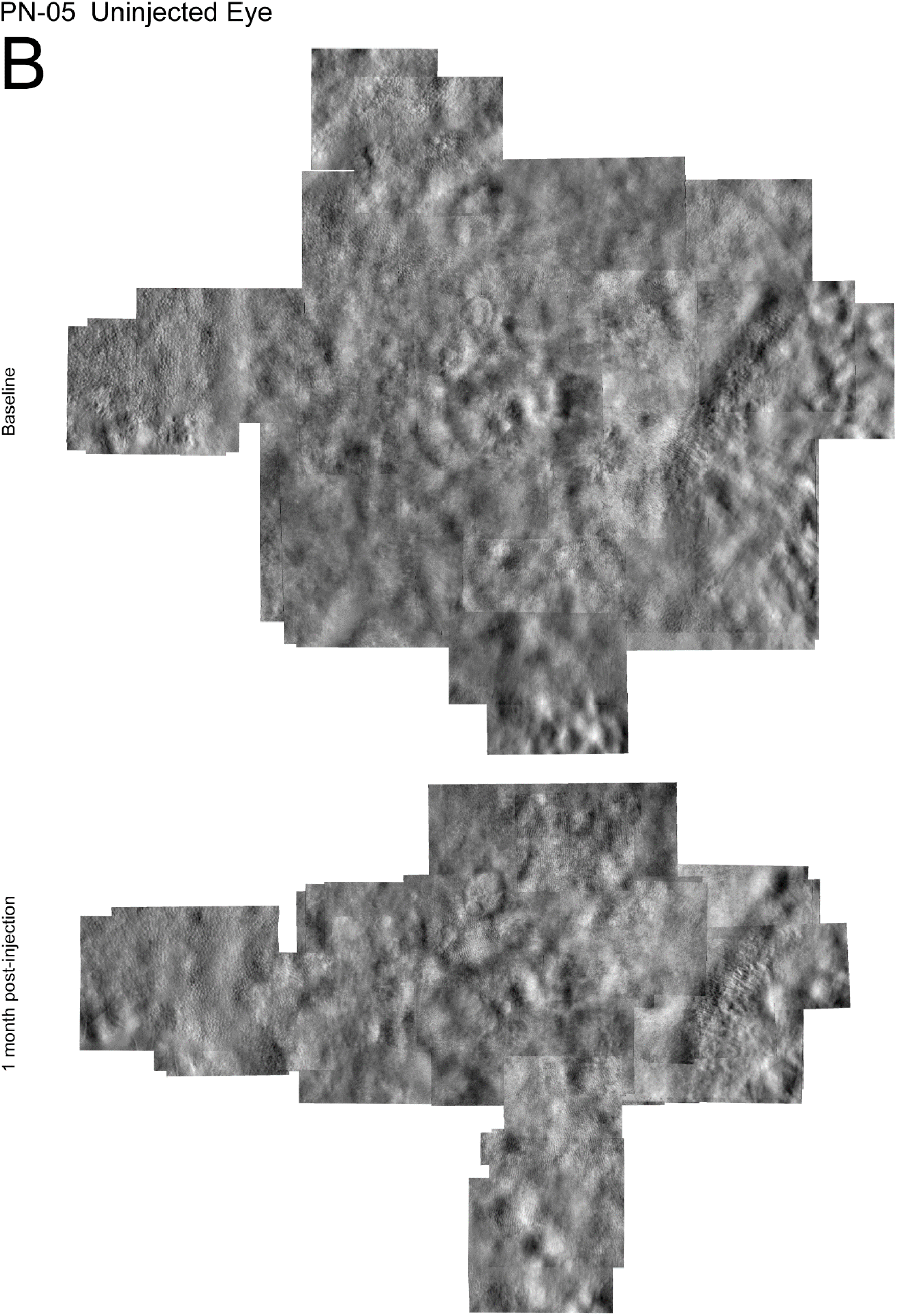

**Supplemental Figure 5A.**
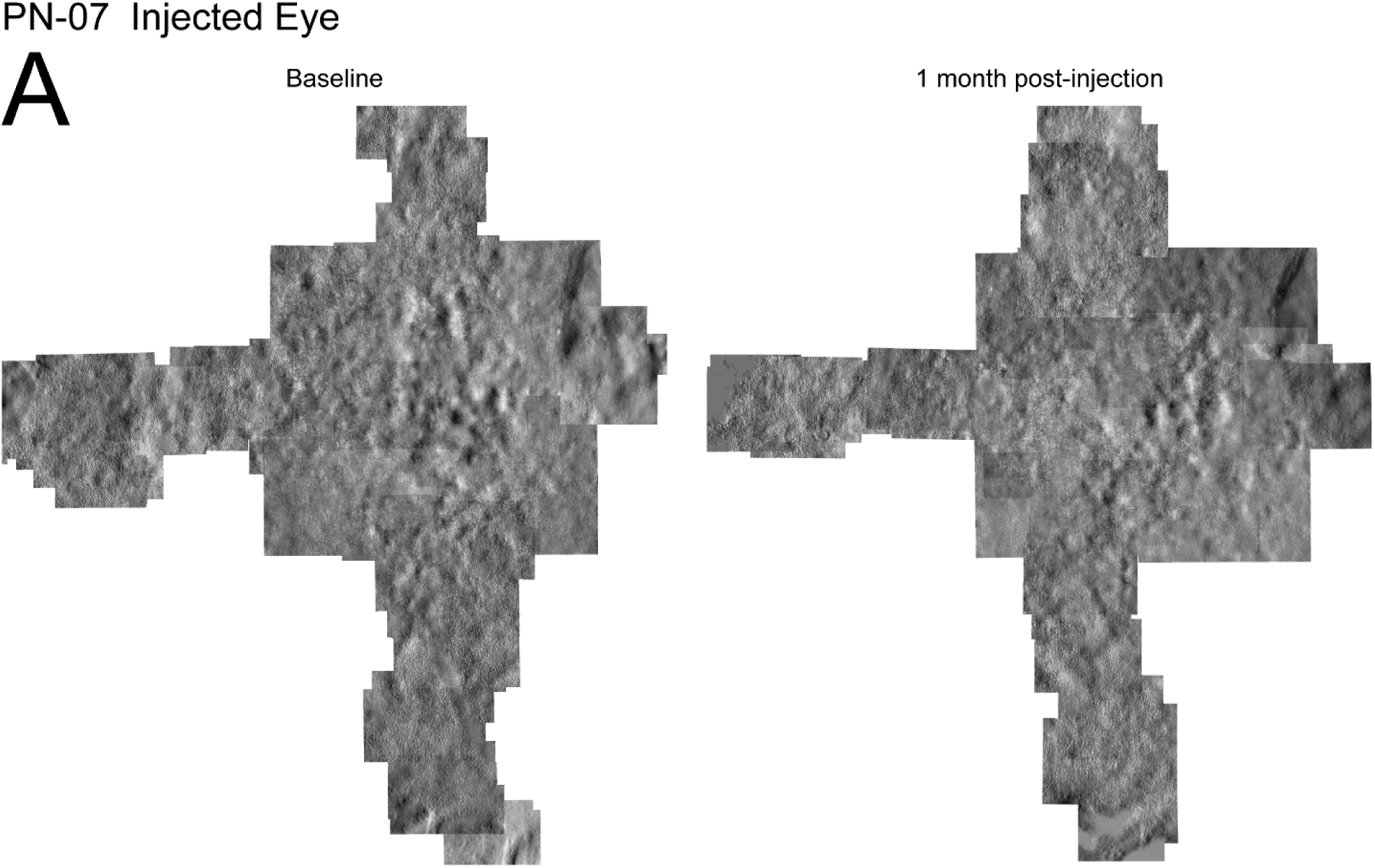

**Supplemental Figure 5B.**
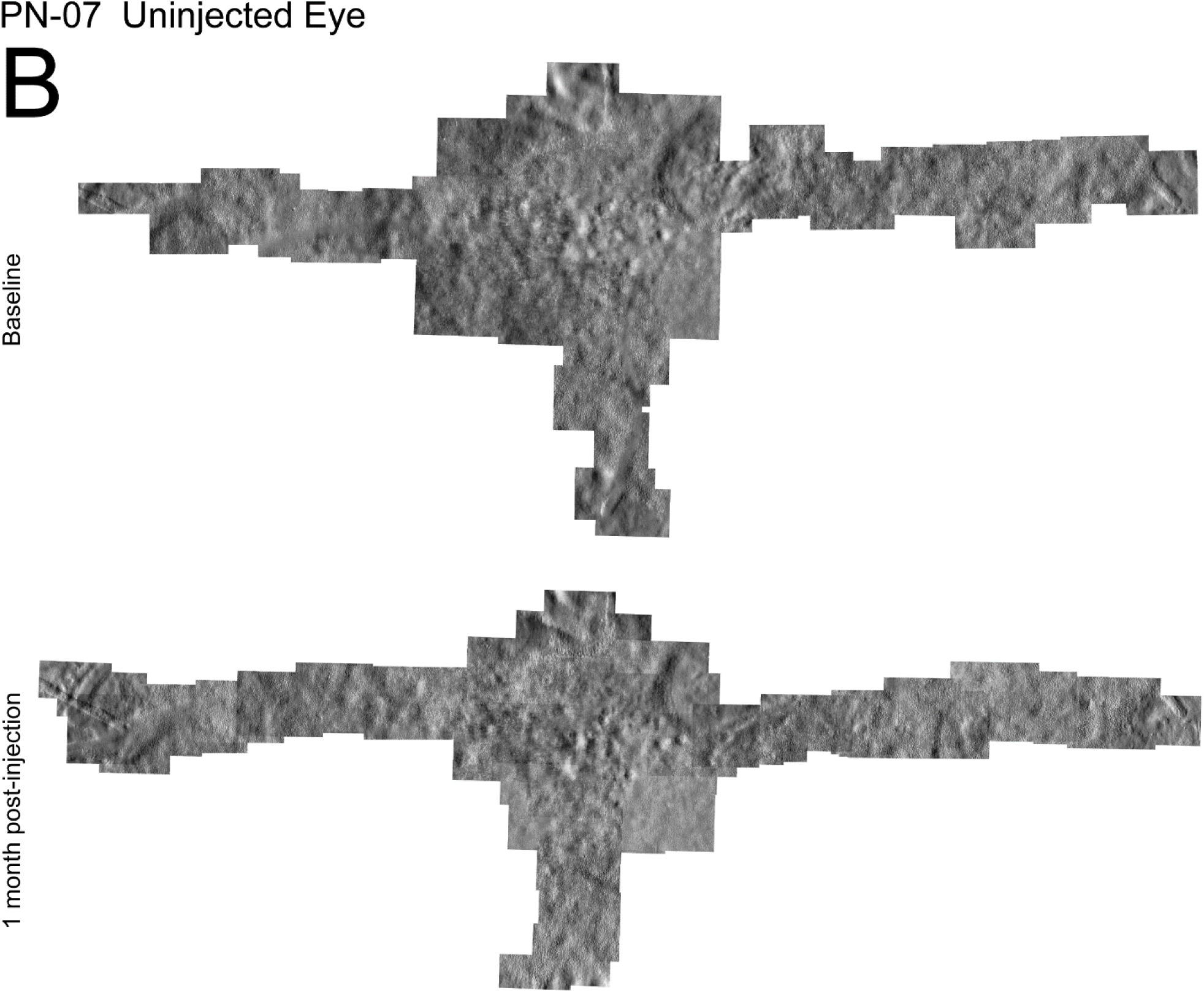

**Supplemental Figure 6A.**
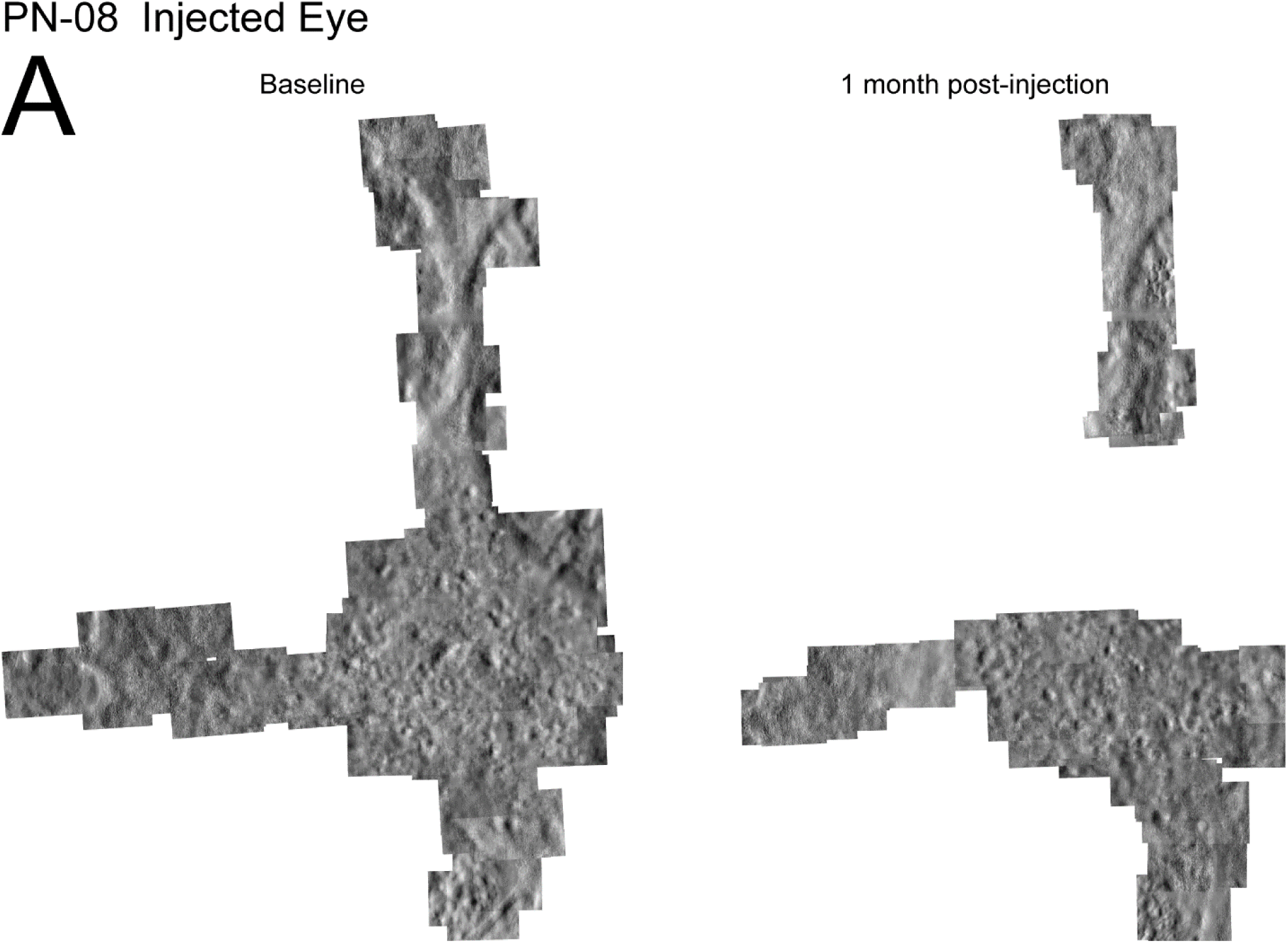

**Supplemental Figure 6B.**
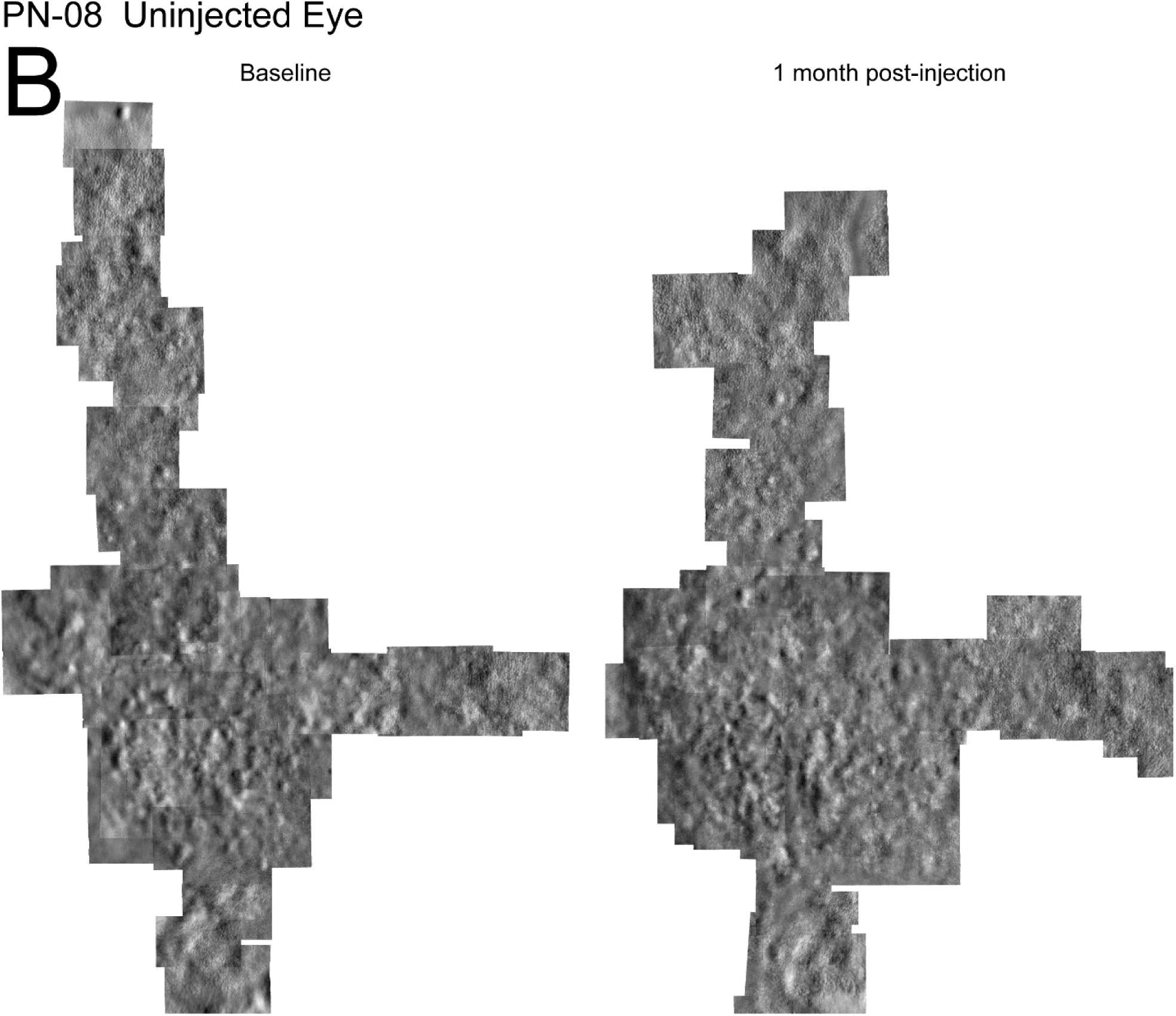

**Supplemental Figure 7A.**
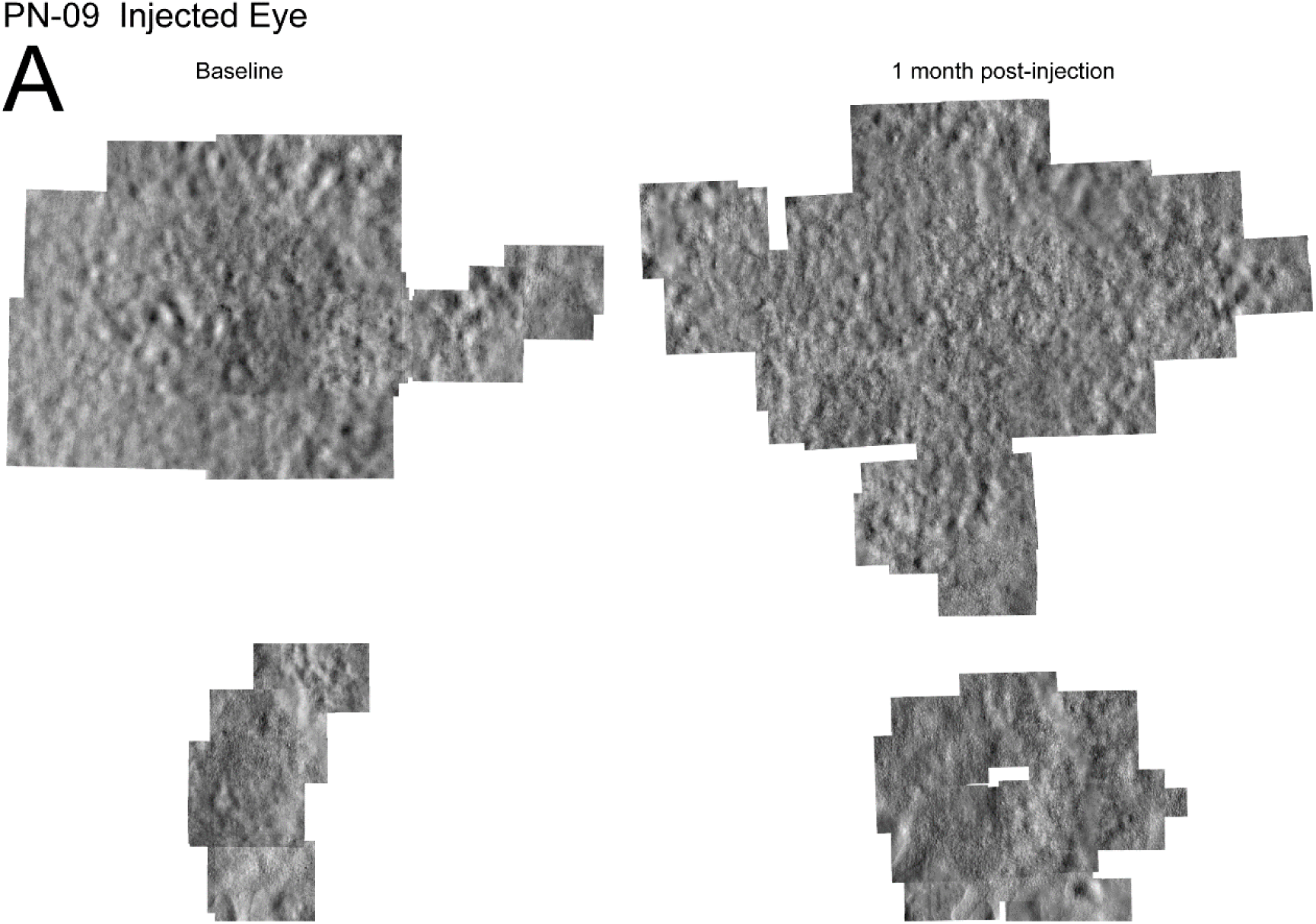

**Supplemental Figure 7B.**
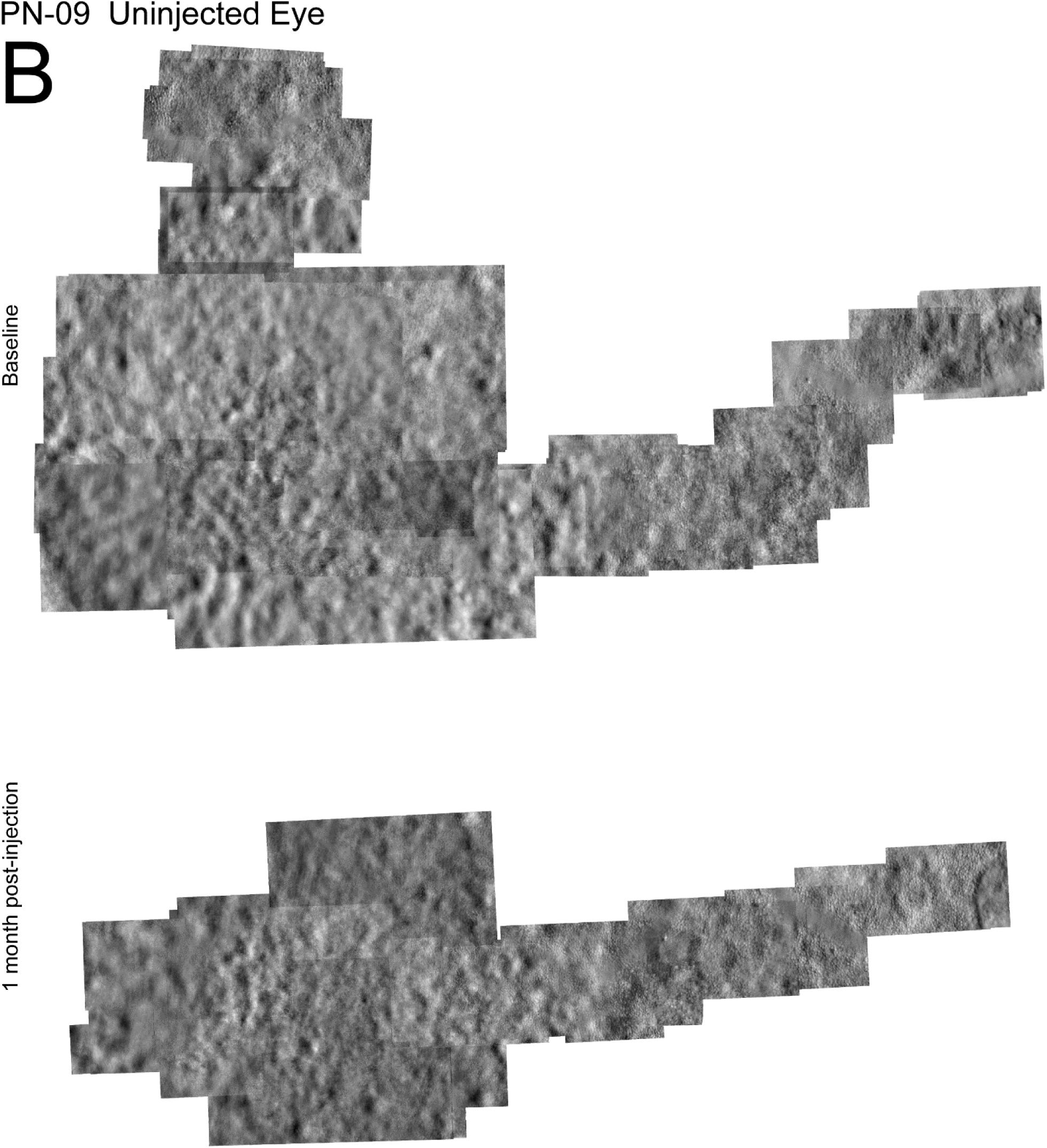

**Supplemental Figure 8A.**
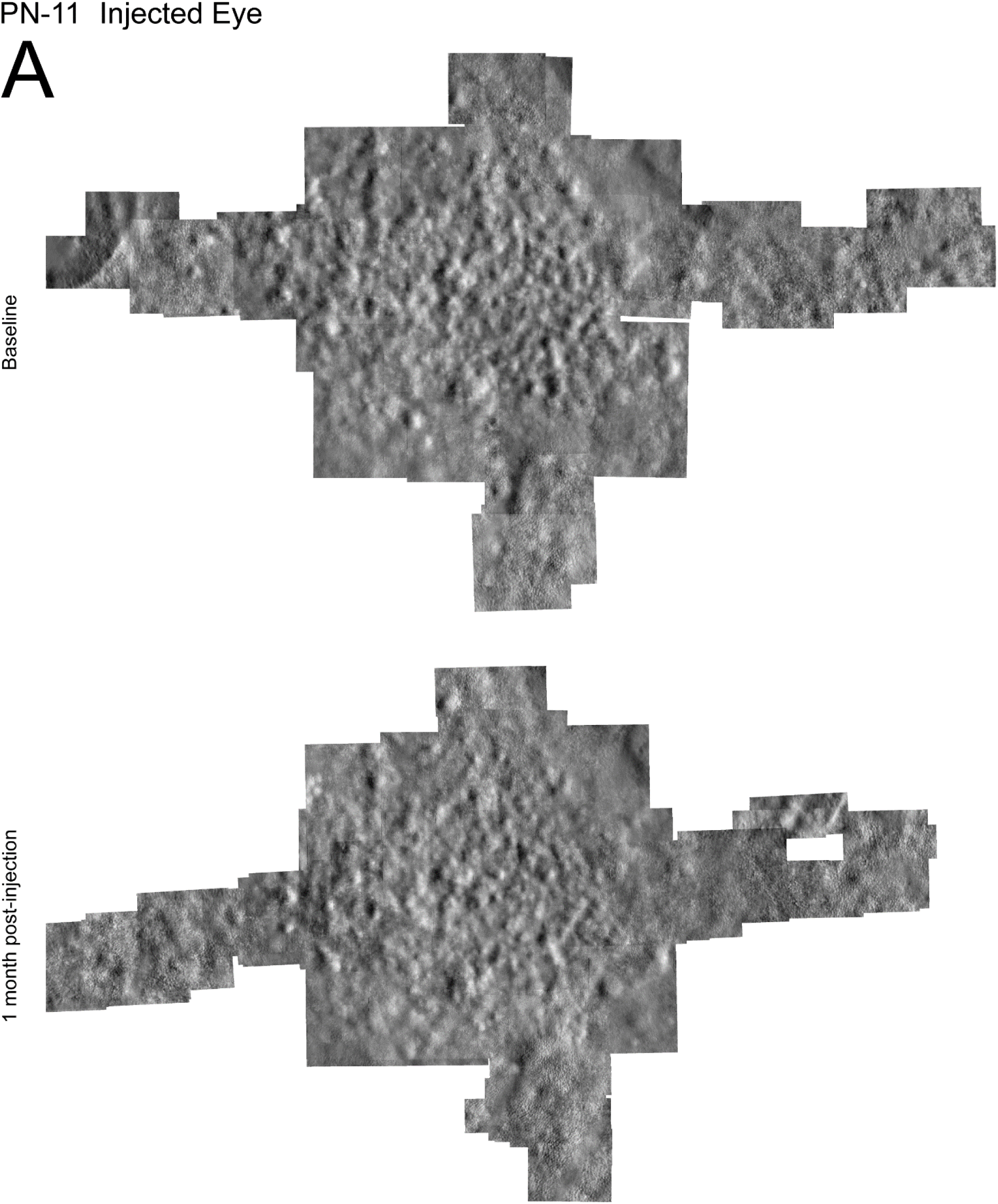

**Supplemental Figure 8B.**
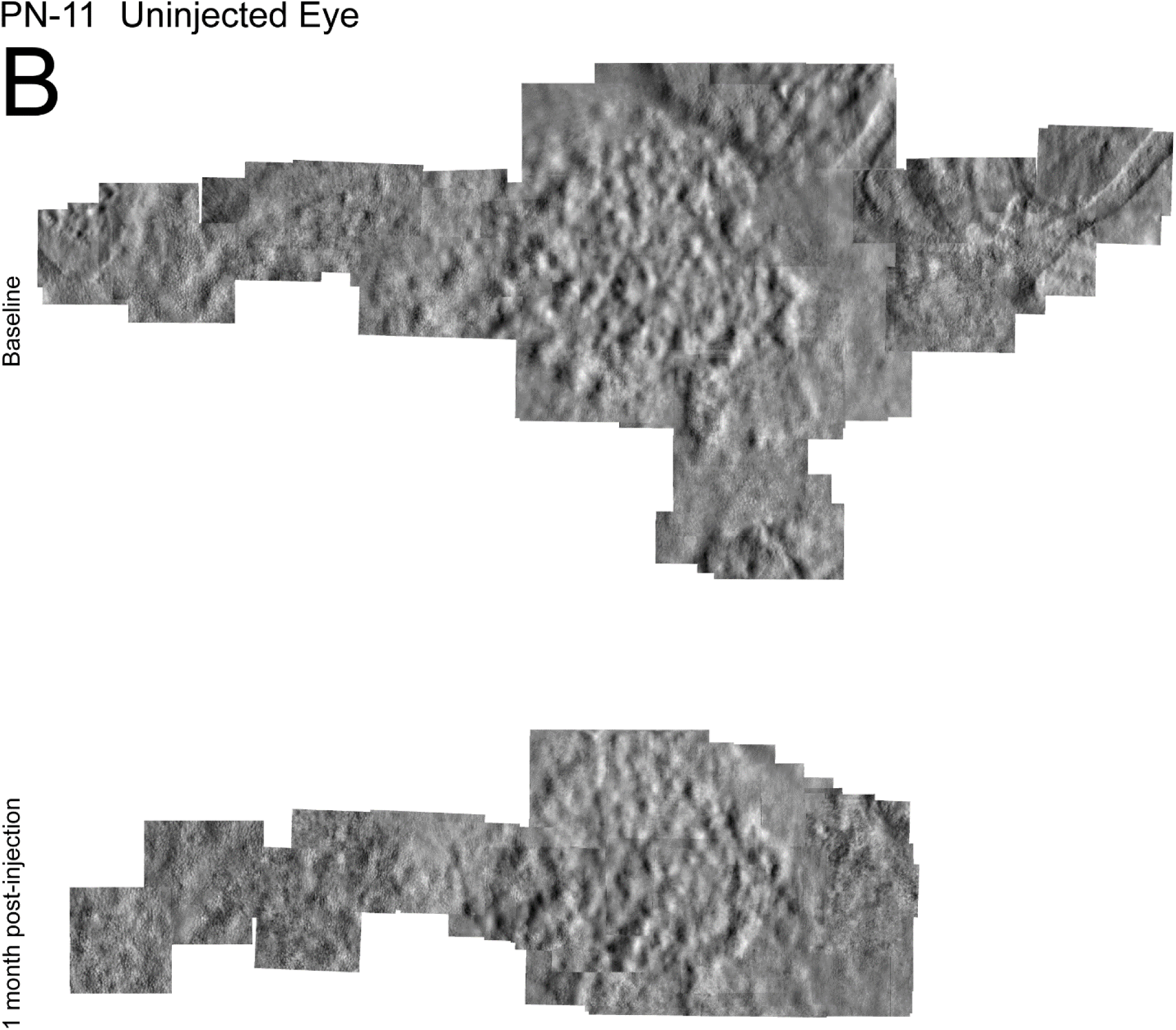

Supplemental Figures 9-17: All AO ROIs used for cone identification and cone density measurements in injected and uninjected eyes. **9:** PN-01 **10:** PN-03 **11:** PN-04 **12:** PN-05 **13:** PN-06 **14:** PN-07 **15:** PN-08 **16:** PN-09 **17:** PN-11 **Top row:** ROIs showing the cone mosaic at baseline in the injected eye. **Second row:** ROIs showing the cone mosaic at one-month post-injection in the injected eye aligned to the baseline ROIs. **Third row:** ROIs showing the cone mosaic at baseline in the uninjected eye. **Bottom row:** ROIs showing the cone mosaic at one-month post-injection in the uninjected eye aligned to the baseline ROIs.

**Supplemental Figure 9.**
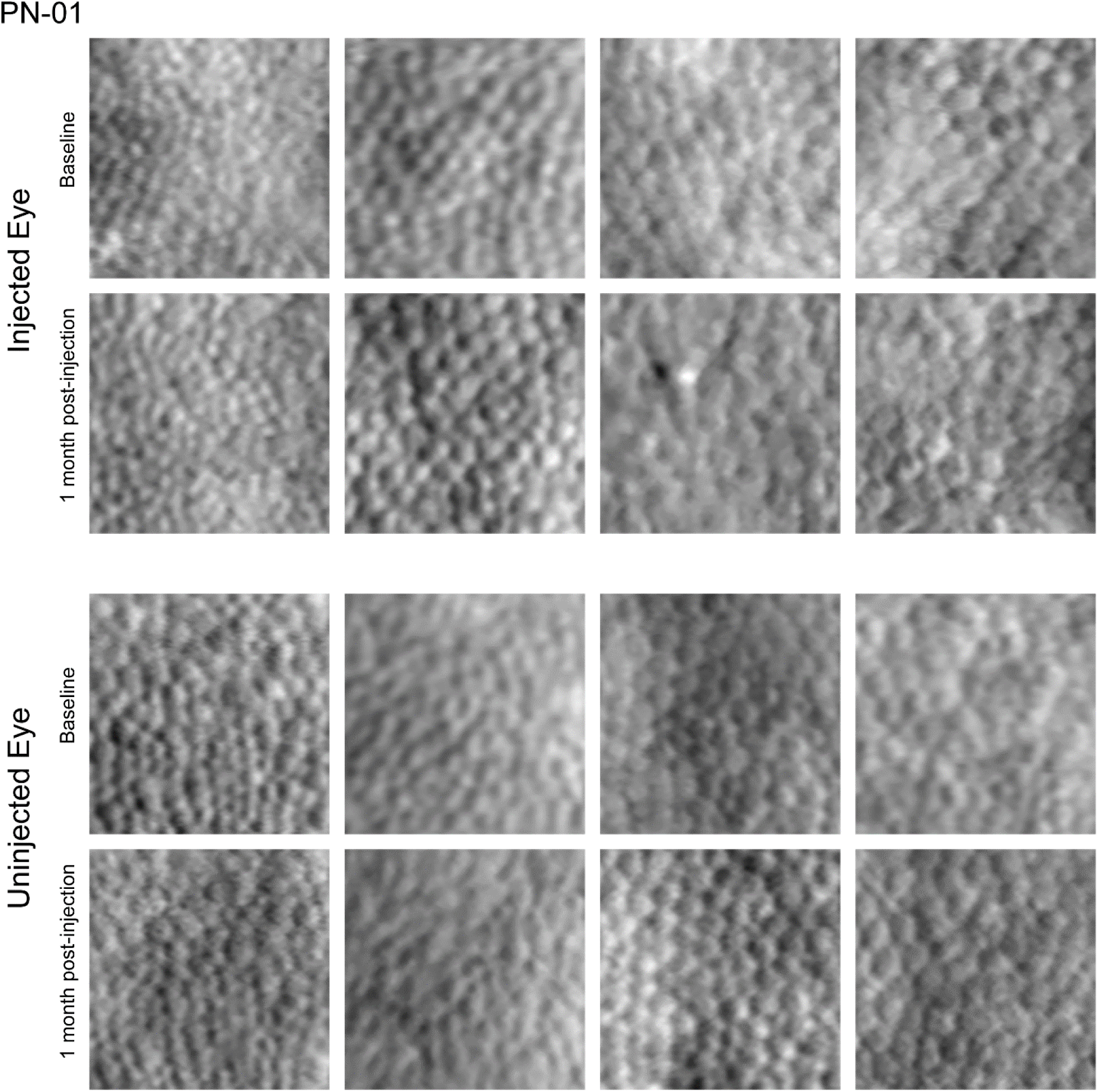

**Supplemental Figure 10.**
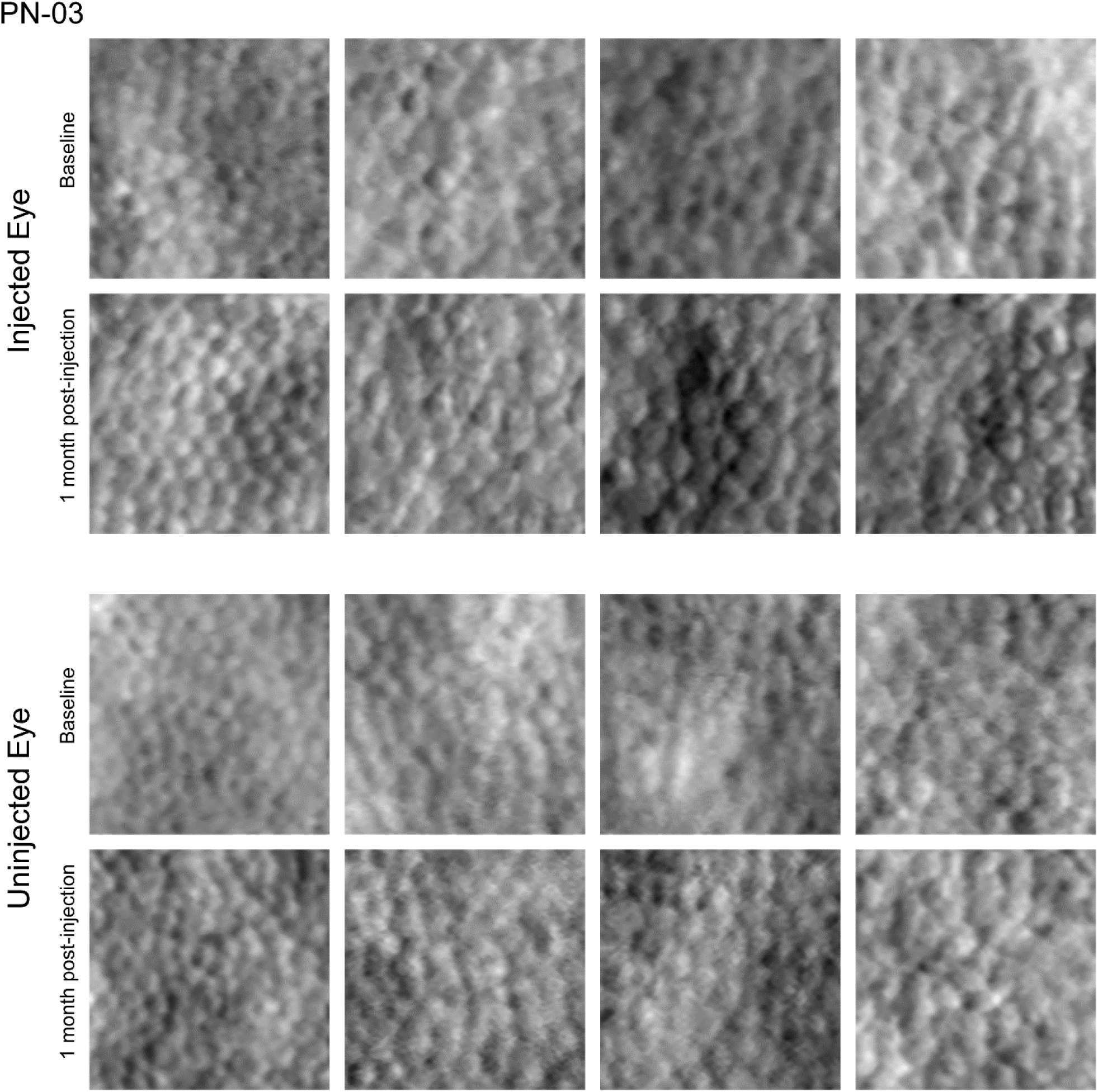

**Supplemental Figure 11.**
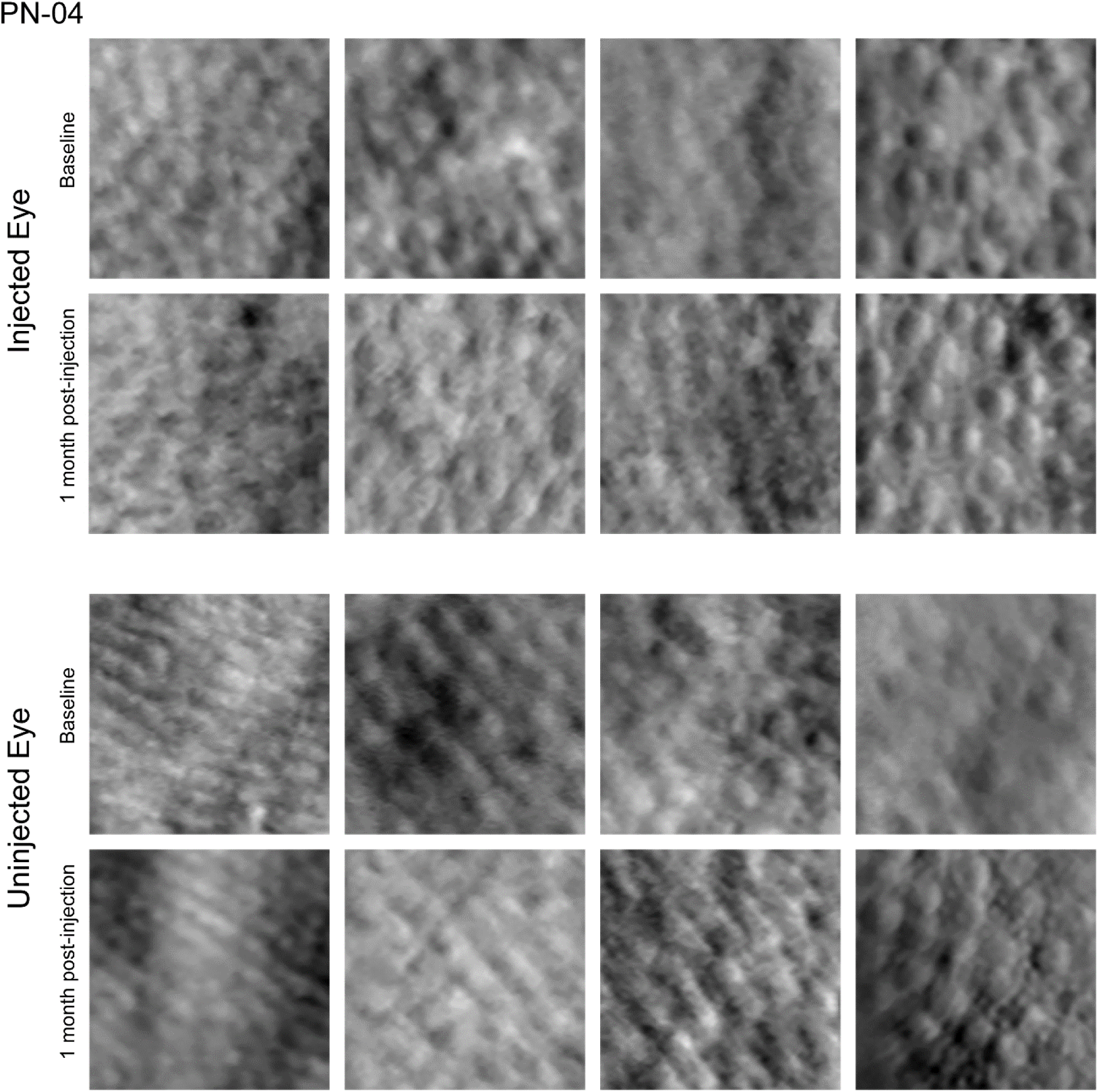

**Supplemental Figure 12.**
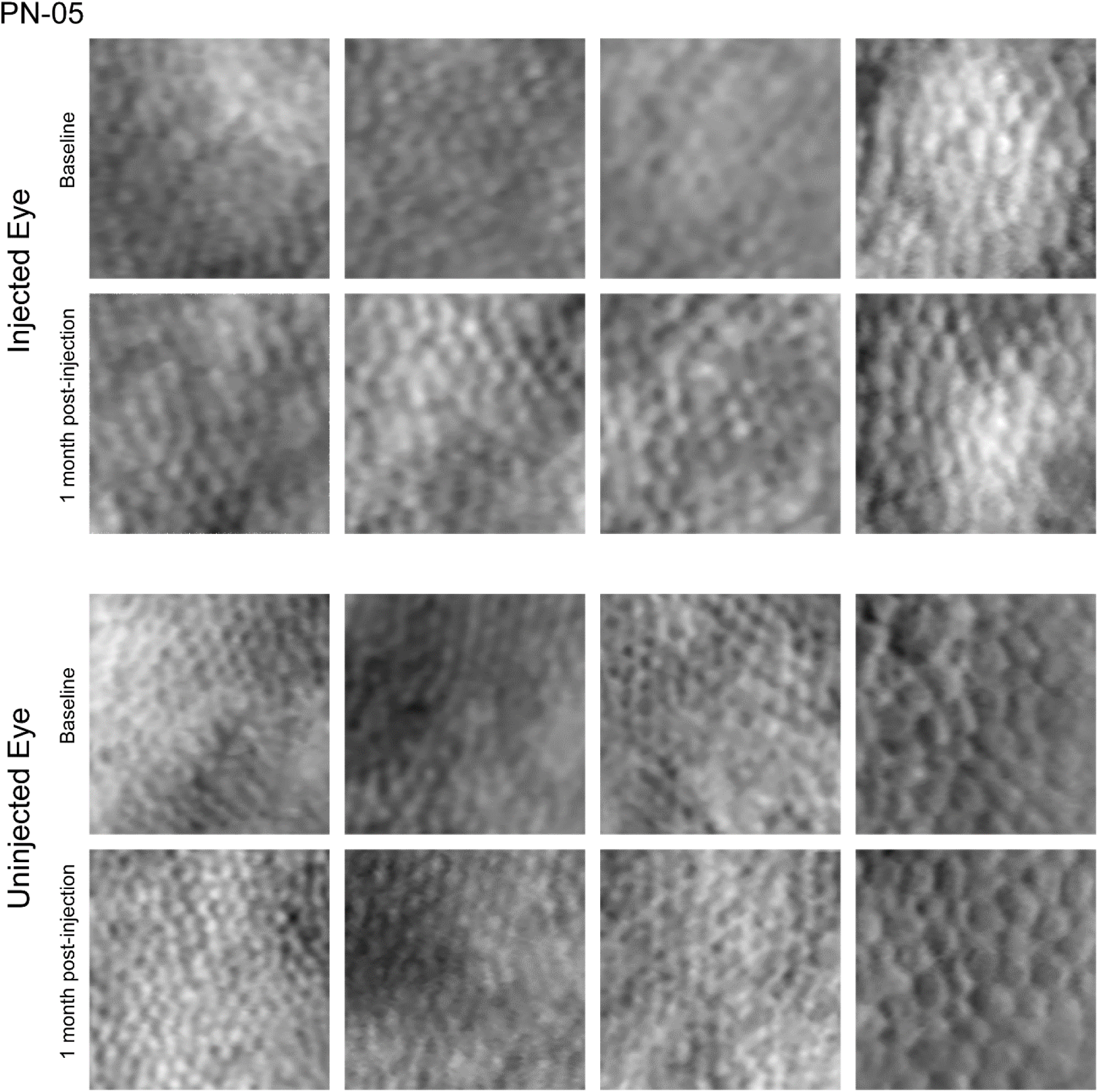

**Supplemental Figure 13.**
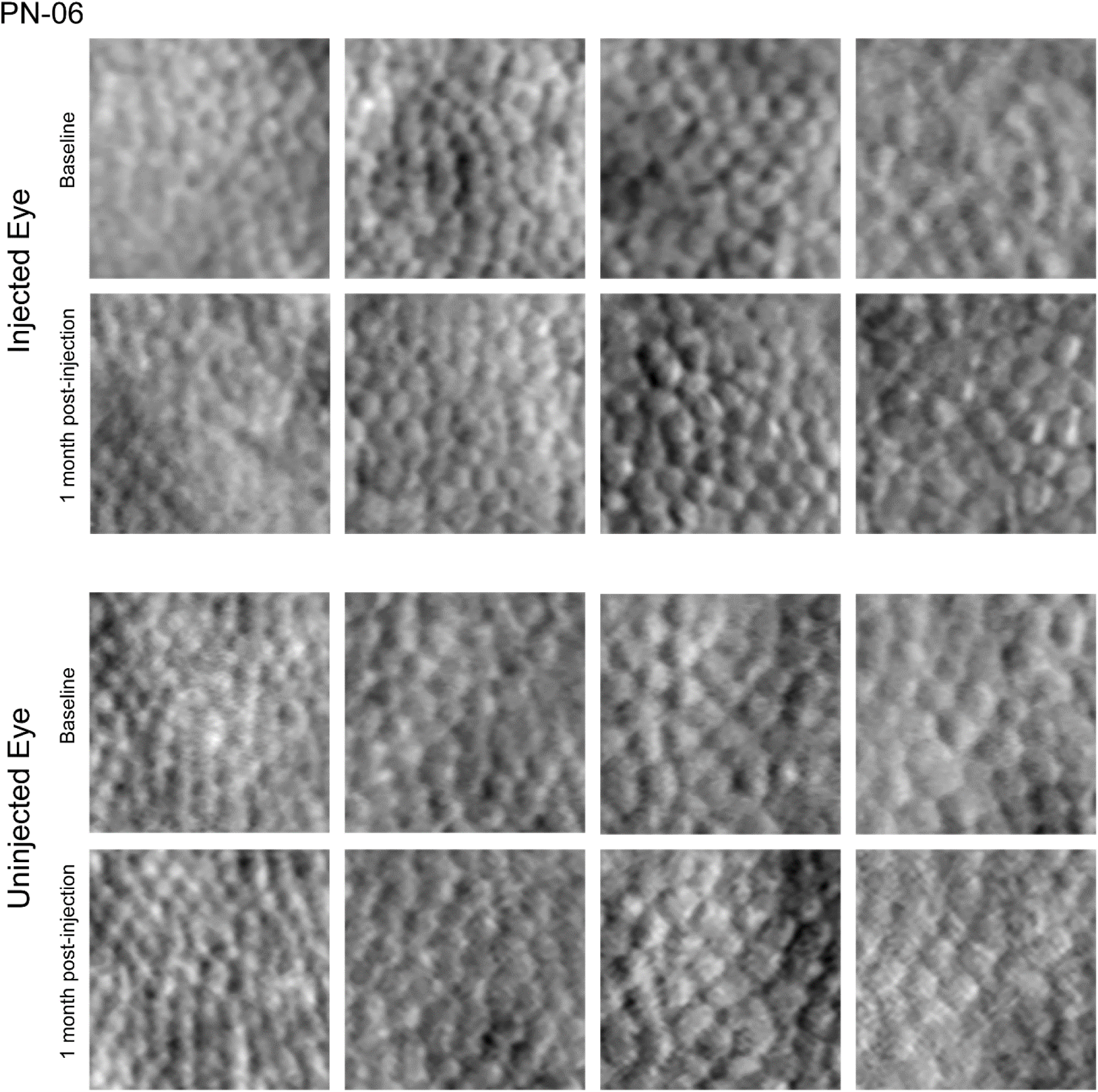

**Supplemental Figure 14.**
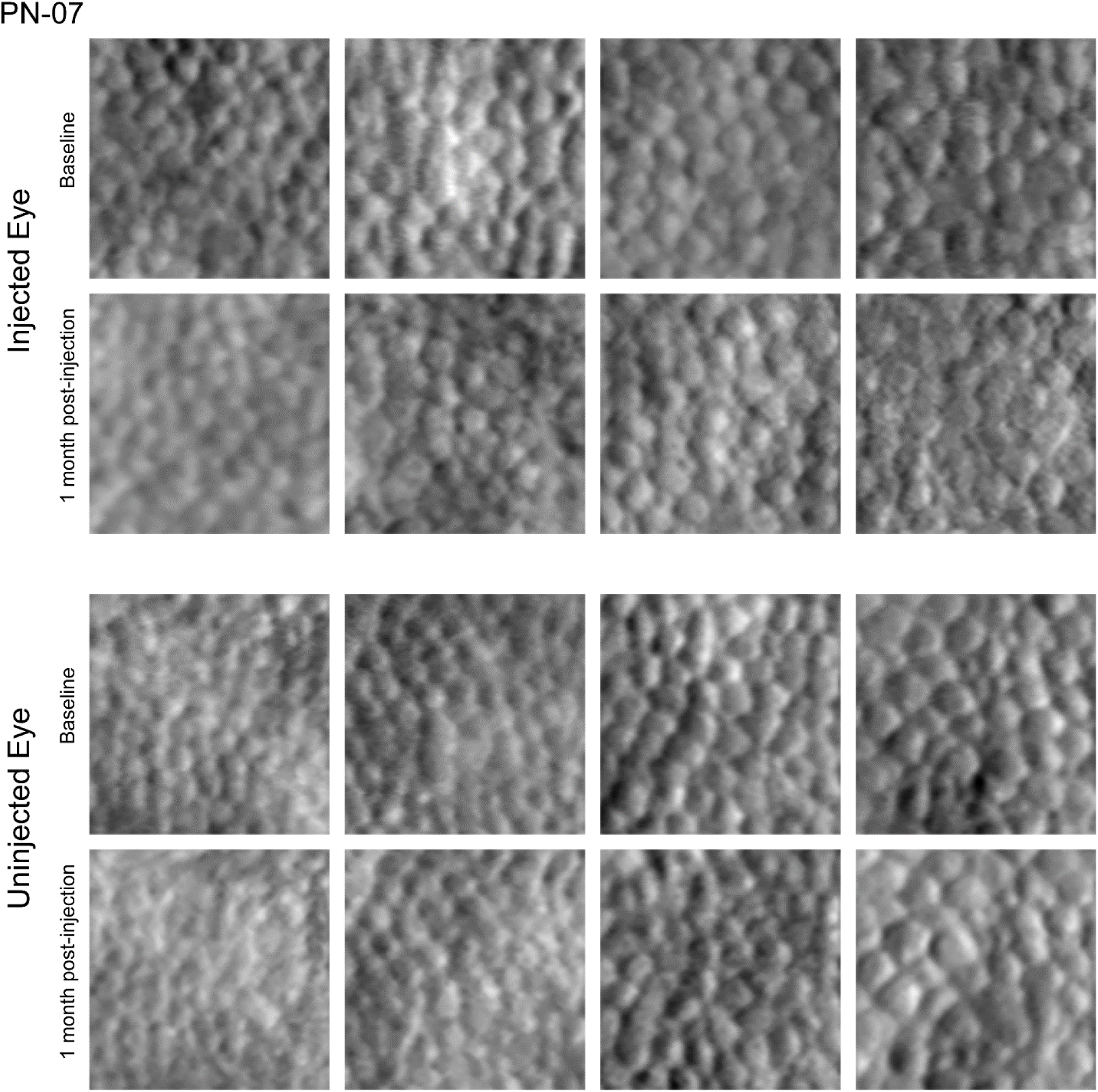

**Supplemental Figure 15.**
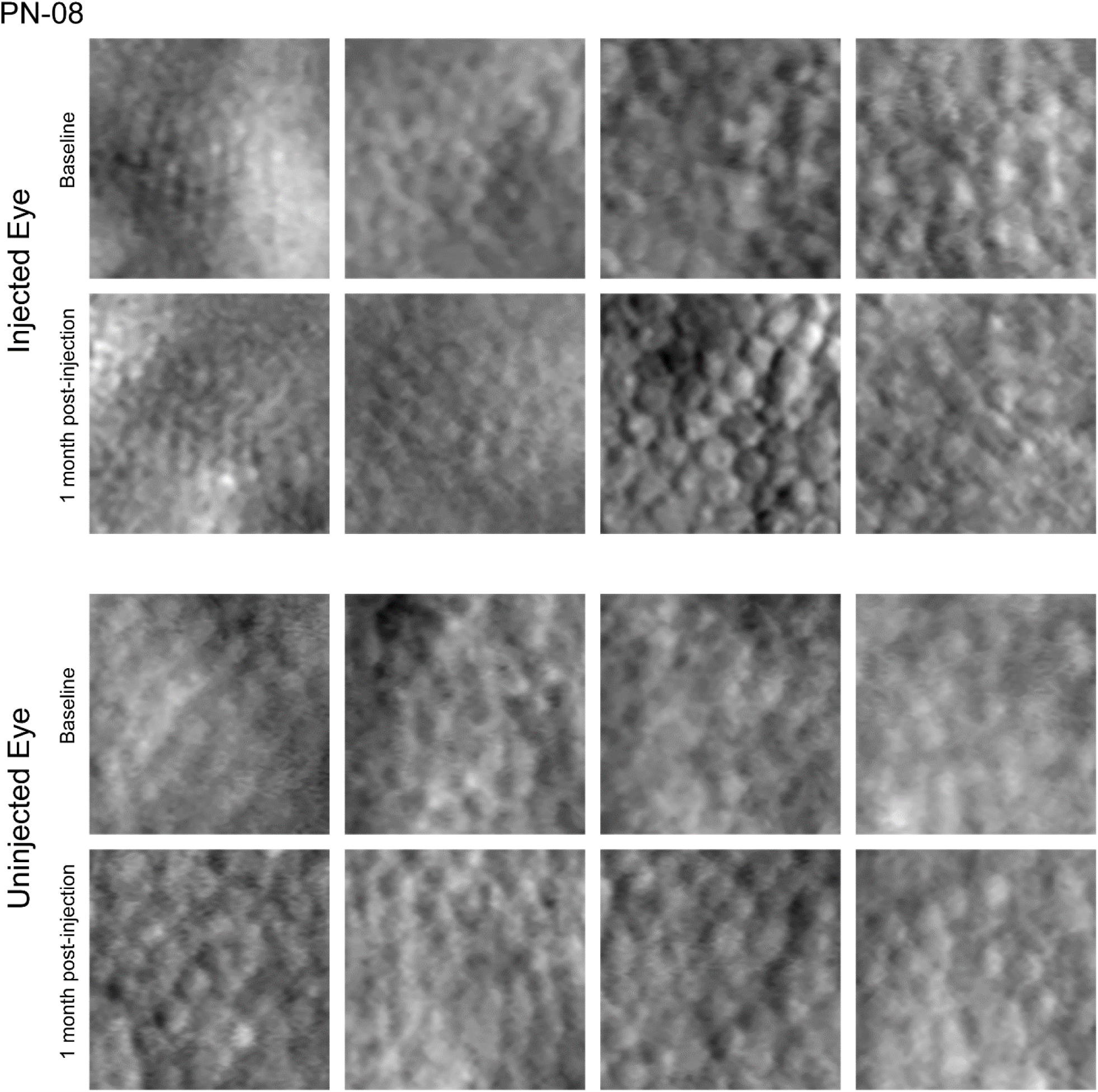

**Supplemental Figure 16.**
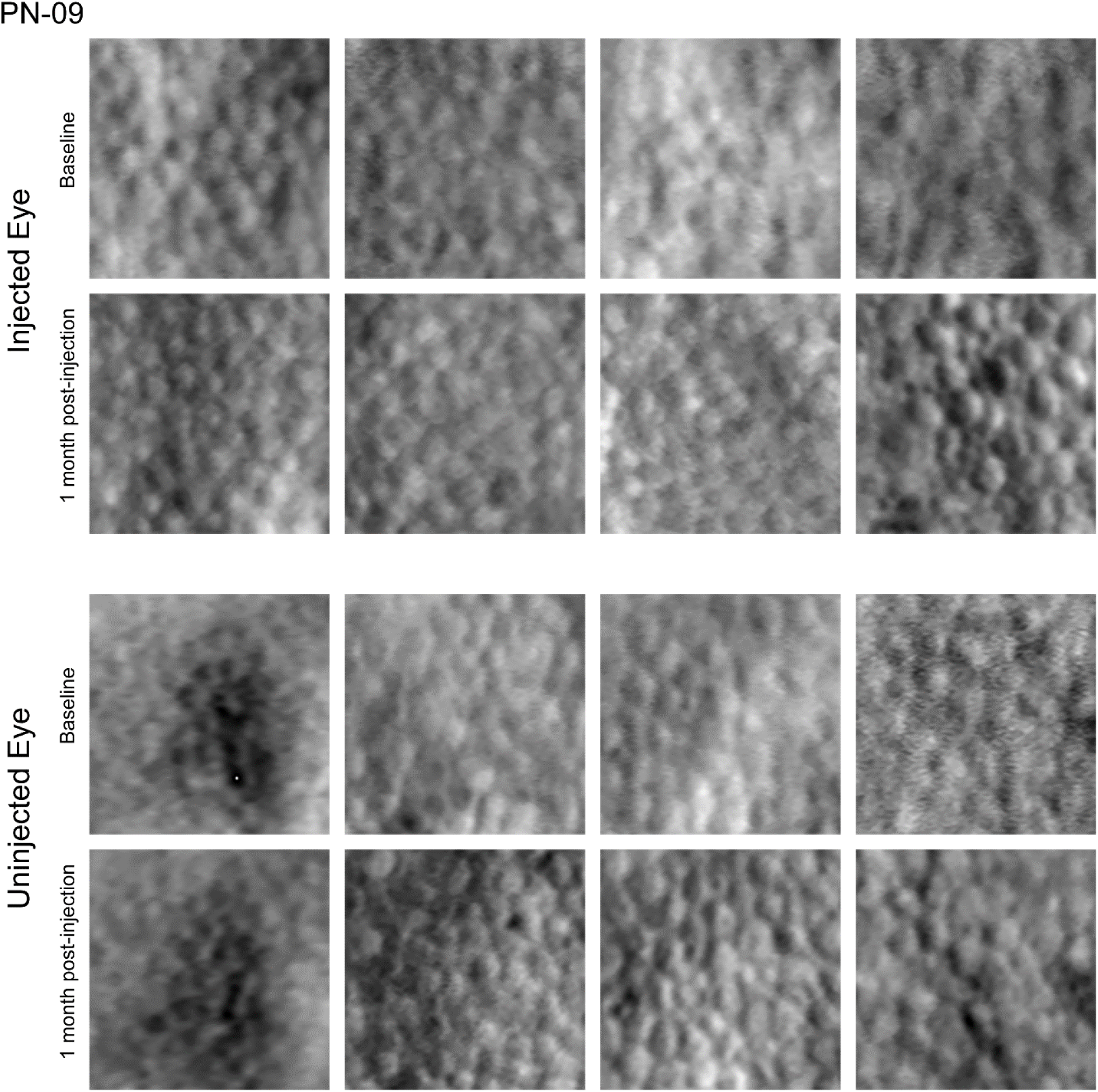

**Supplemental Figure 17.**
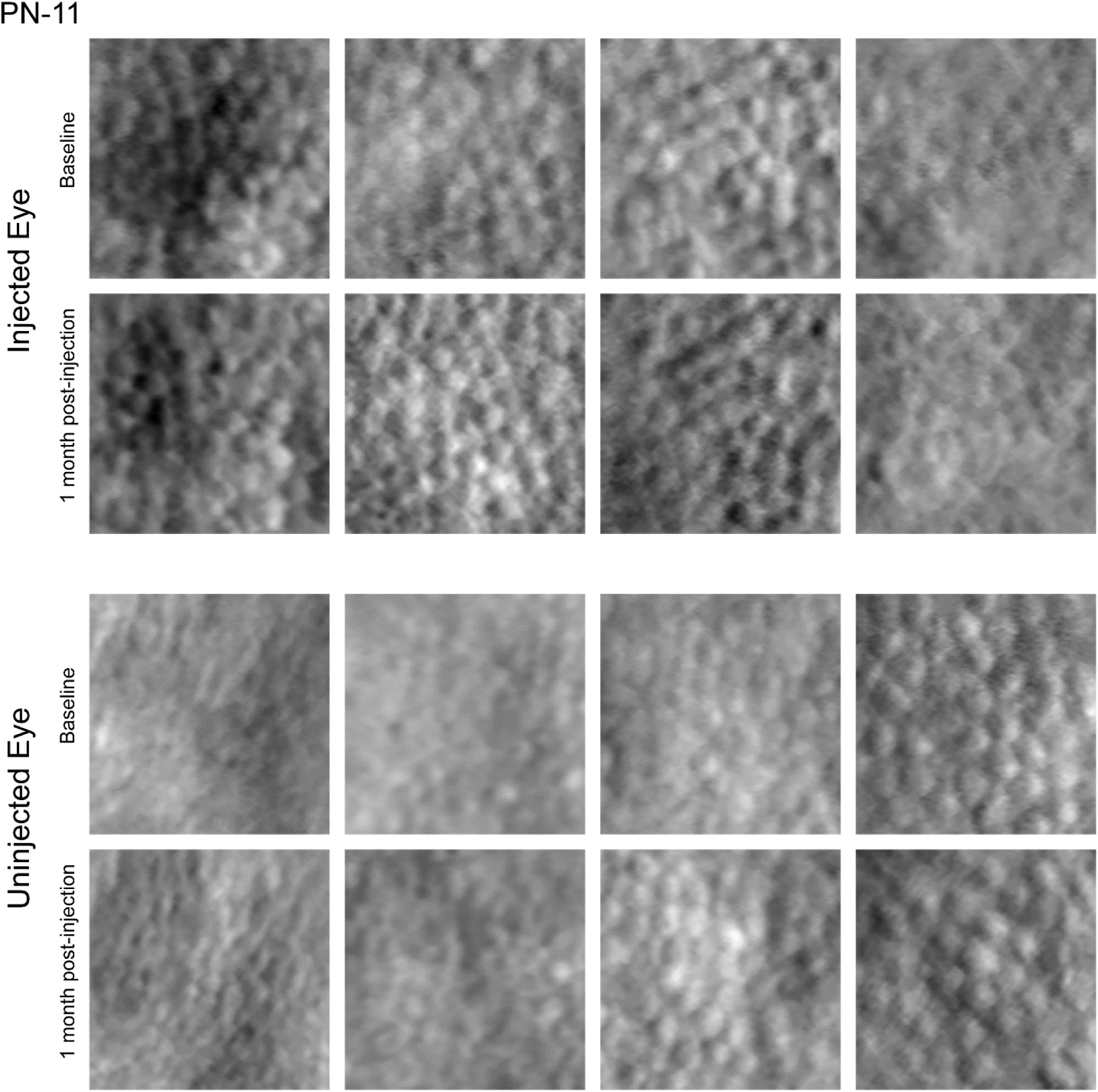

